# A genealogical estimate of genetic relationships

**DOI:** 10.1101/2021.08.18.456747

**Authors:** Caoqi Fan, Nicholas Mancuso, Charleston W.K. Chiang

## Abstract

The application of genetic relationships among individuals, characterized by a genetic relationship matrix (GRM), has far-reaching effects in human genetics. However, the current standard to calculate the GRM generally does not take advantage of linkage information and does not reflect the underlying genealogical history of the study sample. Here, we propose a coalescent-informed framework to infer the expected relatedness between pairs of individuals given an ancestral recombination graph (ARG) of the sample. Through extensive simulations we show that the eGRM is an unbiased estimate of latent pairwise genome-wide relatedness and is robust when computed using genealogies inferred from incomplete genetic data. As a result, the eGRM better captures the structure of a population than the canonical GRM, even when using the same genetic information. More importantly, our framework allows a principled approach to estimate the eGRM at different time depths of the ARG, thereby revealing the time-varying nature of population structure in a sample. When applied to genotyping data from a population sample from Northern and Eastern Finland, we find that clustering analysis using the eGRM reveals population structure driven by subpopulations that would not be apparent using the canonical GRM, and that temporally the population model is consistent with recent divergence and expansion. Taken together, our proposed eGRM provides a robust tree-centric estimate of relatedness with wide application to genetic studies.

## Introduction

Genetic relationships among individuals, commonly characterized by a genetic relationship matrix (GRM), has fueled major advances in modern human genetics. Its applications include the detection of population structure^1,2^, adjusting for shared genetic backgrounds in genome-wide association testing^3–10^, and heritability estimation^11^. Historically, genetic relationships across pairs of individuals in a known pedigree were estimated using the expected proportion of co-inherited alleles, which neglects the variance in the distribution of alleles from meiosis^12–14^. The advent of high-throughput genomics has enabled estimating pairwise relationships directly from genotype data, without the need to rely on expectations determined from an inheritance model^15^.

The current standard to calculate the GRM is based on computing a weighted expectation across genotyped variants (i.e. identity-by-state or IBS)^11,12,14^. While straightforward to compute, this approach generally does not utilize linkage information between markers (though also see ref. ^4,16^). The canonical GRM is not designed to reflect the shared genealogies that connect everyone in a population and inadequately reflects the contribution of ungenotyped variation to relatedness^17–19^. Thus, genome-wide IBS-based relatedness is sensitive to ascertainment biases of genetic variation and only partially captures individuals’ relationships compared with relatedness based on the underlying genealogies of the population. An identity-by-descent (IBD) based GRM could incorporate linkage information to infer finer-scale genetic relationships underlying the structure or demographic history of the study population. However, current bioinformatic approaches estimating shared IBD segments are subject to technical and methodological constraints that effectively limit the resolution of inferred relatedness to only the most recent branches nearing the tips of the underlying genealogies (i.e. over the last 50-100 generations)^12,13,20–24^. Because of its methodological simplicity, the canonical, IBS-based, GRM continues to be the standard in statistical genetics despite its shortcomings^12,20^. Nevertheless, these shortcomings motivate the search of an approach that better captures the genealogical relatedness in a population sample.

In this study, we describe a model for pairwise relatedness using a coalescent-based framework relating everyone in a population sample. Given a coalescent tree at a locus, we define relatedness between individuals by tracing the tree backwards to a single common ancestor. The locus-specific tree provides generalized IBD information across the population sample, unlike conventional definitions of IBD that are restricted to recent branches of the tree in forms of detectable IBD segments of defined multi-generational pedigrees. The entire genealogy of a sample of individuals can be represented by a sequence of coalescent trees, encoded in a structure called the Ancestral Recombination Graph^25,26^ (ARG). The ARG carries substantial linkage information as mutations on the same branch are by definition linked, and historical recombination events are encoded across the sequence of trees. In practice, the ARG is inferred through haplotypic linkage that exists in genetic data. As such, a genealogical measure of relatedness conditioned on the ARG can exploit linkage information that is commonly ignored in the canonical GRM.

Here, we propose a novel coalescent-based framework to estimate the expected genetic relatedness, or the eGRM, for pairs of individuals given the ARG of the population. Conceptually, the eGRM is based on the expected number of mutations occurring randomly on each branch of the ARG, rather than directly genotyped variants. Our framework provides two primary benefits compared with previous approaches. First, because the ARG encodes historical recombination events and the estimation of the ARG generally leverages patterns of haplotype sharing, the eGRM in practice is expected to be more robust to ungenotyped genetic variation and retains greater information of IBD relatedness among individuals than the canonical GRM. Second, and more importantly, our framework seamlessly provides insights to the time-varying nature of population structure by estimating relatedness at specific depths in the coalescent tree. To enable efficient calculation of the eGRM, our framework leverages recent computational advancements for scalable ARG inferences^27,28^, thus enabling investigation of populations in larger datasets.

We characterized the behavior of our ARG-based eGRM through extensive simulations starting from standard population genetic models. In simulations of a single, exponentially growing, population, we demonstrate that the eGRM better captures latent genome-wide relatedness compared with the canonical GRM. Importantly, we find the improved performance of eGRM is robust when performing inference using noisier ARGs inferred from a subset of common genotyped variants rather than true ARGs. It is believed that common variants are not sufficiently informative to detect recent population structure^29^. However, in simulations of a recently structured population with multiple demes, we find that principal component analysis (PCA) of the eGRM better reflects overall population structure and more accurately identifies each deme, compared to PCA of the canonical GRM. Finally, in analyses of 2,644 genotyped samples from Northern and Eastern Finland^21^, we observe that PCs derived from the eGRM reveal fine-scale structure previously not identified when using the canonical GRM. We estimate multiple partial eGRMs at multiple epochs across the history of the sample and show that time-specific patterns of population structure are qualitatively similar to simulated results of a recently structured population model, which is consistent with the known history of this region of Finland.

## Results

### Method Overview: a conceptual shift in defining genetic relatedness

The eGRM, conditioned on the ARG, is conceptually different from the canonical GRM. We demonstrate this difference through a toy example on a single genealogical tree with 4 branches and 6 mutations (**Figure 1A**). The canonical GRM is variant-centric and is the average of the six relatedness matrices based on each mutation. The eGRM, however, defines relatedness through tree branches that relate a pair of haplotypes. Assuming constant mutation rates across branches, the eGRM is the average of the four relatedness matrices based on each branch, weighted by their branch lengths. A single tree is shown in **Figure 1A** for simplicity, but the eGRM can be generalized to a sequence of trees along a chromosome by weighting each tree by its total branch length times the number of base pairs covered by each tree (**Methods**). In this toy example, haplotypes *a* and *b* are expected to be equally related to *c* in the eGRM, while in the canonical GRM *b* will be more closely related to *c*. Under the canonical GRM framework, the relative genetic distance to *c* is subject to the randomness and ascertainment of mutations. Instead of relying on ascertained mutations, branch lengths from the true ARG (or from the inferred ARG, reconstructed based on linkage information among nearby markers) provide an estimate of genetic relatedness that is more robust to ascertainment effects. In addition, while the eGRM is defined as a function of the genealogy, it maintains the mathematical properties of canonical GRMs (e.g., positive definiteness) as eGRM is the expectation of the canonical GRM. The eGRM is thus compatible with all downstream applications of the GRM. To help distinguish between various eGRM estimators, here we define some useful notation. We denote the eGRM estimated conditioned on the true ARG as **EK**. When conditioned on an ARG inferred from genetic data using either RELATE^28^ or TSINFER+TSDATE^27,30^ we denote such eGRM as **EK**_relate_ and **EK**_tsdate_, respectively. We denote the canonical GRM computed using all latent genetic data as **K**_all_ and using only the observed genetic data as **K**_obs_. In empirical data analysis, **K**_obs_ are constructed using all genotyped data passing quality controls; in simulations, **K**_obs_ are constructed using only 20% of the genetic data oversampled from the common variation of the frequency spectrum (**Methods**) to mimic a genotyping array. Importantly, **EK**_relate_ and **EK**_tsdate_ are constructed only using the same set of the observed genetic data as **K**_obs_.

**Figure 1.**
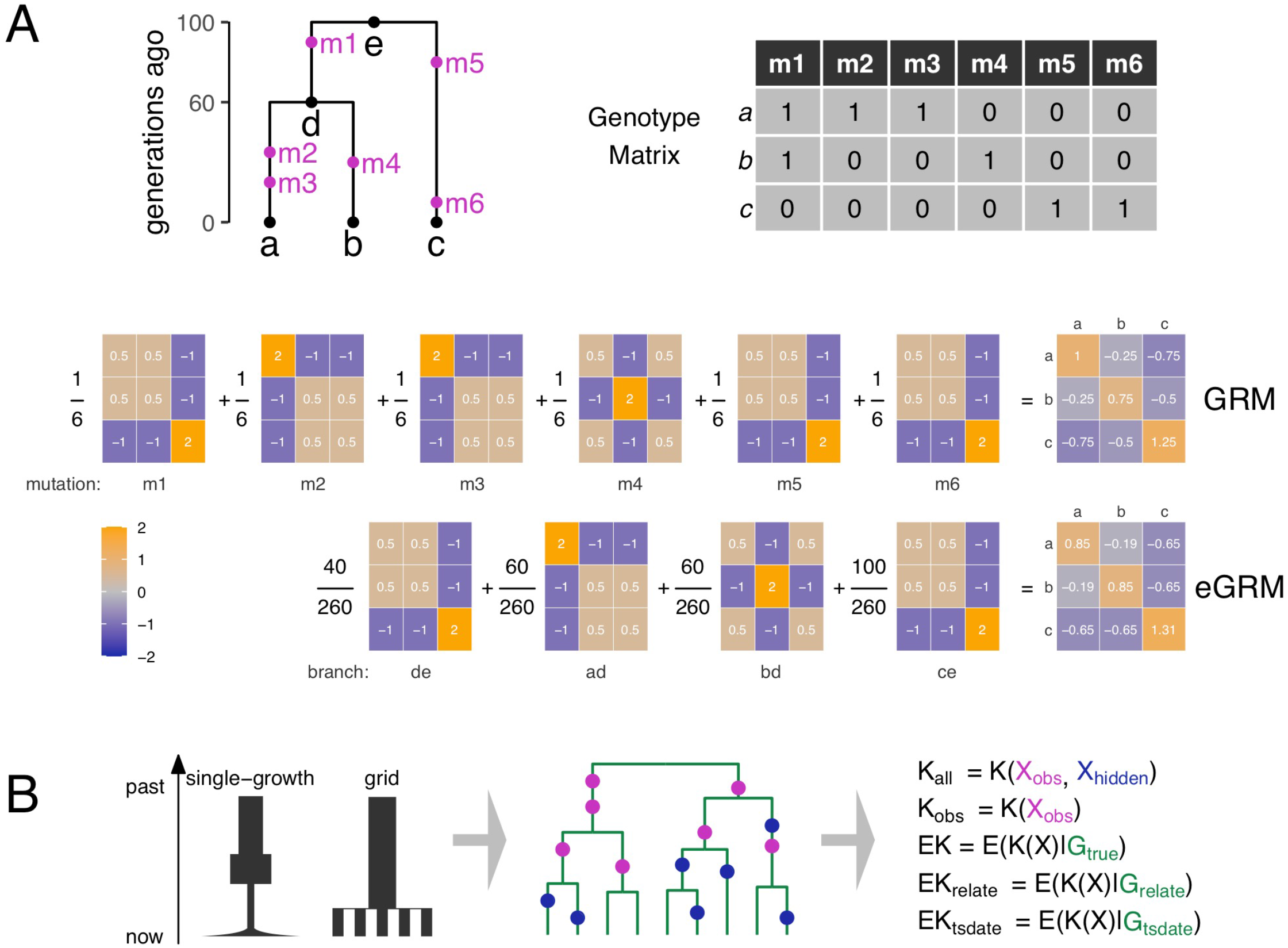
Illustrative example of eGRM and methodological overview. **(A**) An illustrative example of a single genealogical tree containing 3 samples, 4 branches and 6 mutations to contrast the eGRM and GRM. Each mutation has a corresponding length-3 haplotype vector (e.g., mutation **m1** has haplotype vector (1, 1, 0)). The “single-variant GRM” can be computed as the outer product of the centered and normalized haplotype vector (e.g.,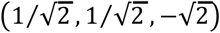 for mutation **m1**) with itself. The canonical GRM is then the unweighted average of the single-variant GRMs of the 6 mutations. The eGRM is based on the 4 branches, weighted by their lengths (i.e., the expected number of mutations on this branch). (**B**) Overview of simulation workflow to test the performance of eGRM. ARGs are simulated by MSPRIME based on a single-growth demographic model and a grid-like spatial structure model. Observed variants are oversampled from common variants to mimic real genotyping array data. We then compute the complete GRM (**K**_all_), the observed GRM (**K**_obs_), eGRM based on the true ARG (**EK**), eGRM based on RELATE- or TSINFER+TSDATE-reconstructed ARG (**EK**_relate_ and **EK**_tsdate_).

### eGRM accurately measures relatedness on a genealogical tree

To establish that the eGRM estimator better reflects genealogical relatedness compared with the canonical GRM approach, we first sought to quantify the performance of eGRM in capturing relatedness in a single tree, defined here as the TMRCA between pairs of individuals, when using the true ARG. We simulated a 1 Mb genetic region with 1,000 individuals under a single population growth model, computed **EK** and **K**_obs_ (see **Methods**; **Figure 1B**). Unsurprisingly, the eGRM based on the true genealogy, **EK**, is better correlated with TMRCA than **K**_obs_ in 97.5% of the simulations (P = 4e-252 by sign test; **Figure 2A**) and more accurately captures recent genetic relatedness between pairs of individuals (**Figure S1A**). More importantly, eGRM constructed using genealogies inferred under RELATE or TSINFER+TSDATE on the same set of observed variants (**EK**_relate_ and **EK**_tsdate_) also showed better correlation with TMRCA than the canonical GRM in ∼70% of the simulations (P < 1e-26 in all cases; **Figure 2A, Figure S1B**), suggesting that the eGRM is robust to noise in inferred ARGs. Our results thus demonstrate a consistent advantage of the eGRM over the canonical GRM in capturing local relatedness represented by TMRCA. Because common variants are individually uninformative for recent relatedness, our results also suggest the eGRM framework based on predominantly common variants can provide insight for the recent part of the genealogical tree.

**Figure 2.**
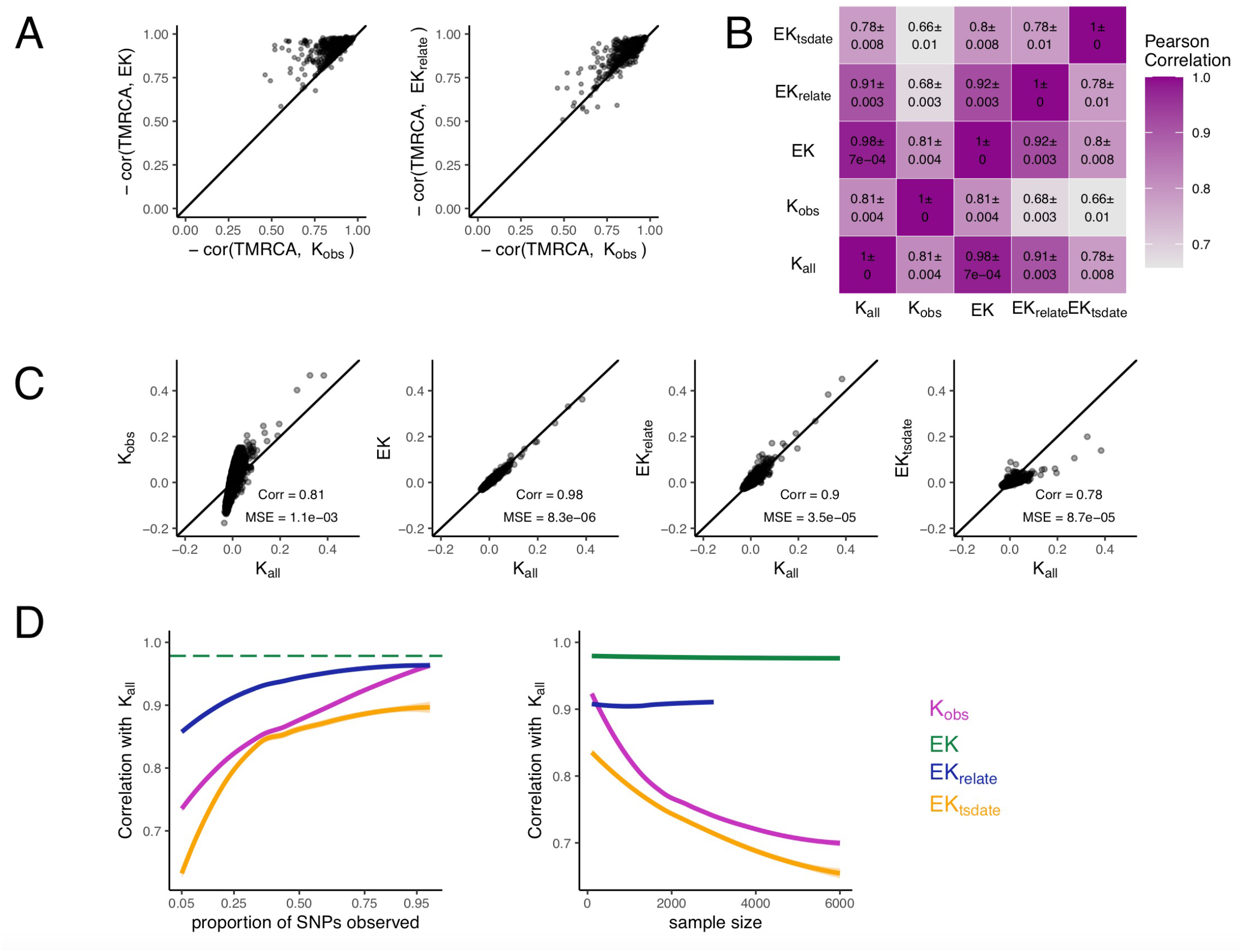
eGRM is highly and unbiasedly correlated with measures of relatedness in simulations. **(A)** Negative Spearman correlation between TMRCA and **K**_obs_, **EK** (left) or **EK**_relate_ (right) on a 1Mb non-recombining locus. Spearman correlation is used because GRM by definition normalizes according to allele frequency to upweight rare mutations, and thus is not expected to correlate linearly with TMRCA. (**B**) Heatmap summarizing the Pearson correlations between GRM and eGRM matrices on a 30Mb chromosome. Note that **EK** and **EK**_relate_ are highly correlated with **K**_all_. (**C**) Scatter plots of the GRM and eGRM values for all pairs of individuals, using the same simulated data as in **B**. All simulations from **A** to **C** simulated 1000 individuals. (**D**) Pearson correlation with **K**_all_, with varying proportion of SNPs observed (sample size is fixed to 1000; left) or varying sample size (20% common SNPs observed; right) on a 30Mb chromosome.

### eGRM provides an unbiased estimate of genome-wide relatedness across realistic scenarios

While TMRCA provides an intuitive measure of the local genetic relatedness between a pair of haplotypes, the eGRM is formulated as the expectation of the latent GRM so that it adheres to the mathematical properties of a GRM necessary for many downstream statistical genetic applications^1–11^. Therefore, we next sought to evaluate how well eGRM is capturing the genome-wide relatedness measured by the latent GRM. Even though the mutation rate in humans is small, we reasoned that a GRM computed using all variants (**K**_all_) in a sufficiently large genomic region and sample will precisely estimate the latent GRM by the law of large numbers. Therefore, we next sought to evaluate how well the eGRM measures genome-wide relatedness, as quantified by the GRM computed from all latent variants (**K**_all_). Briefly, we repeatedly simulated a 30-Mb genomic region of 1,000 individuals with recombination rate set as 1e-8 per bp per generation (see **Methods**). We found that **EK** provides an approximately unbiased estimate of **K**_all_ (Pearson correlation = 0.98 ± 0.0008; **Figure 2B**; regression slope of 0.951, 95% CI [0.949, 0.953], intercept -6.7e-5, 95% CI [-9.1e-5, -4.3e-5]; **Figure 2C**), when compared with **K**_obs_ (Pearson correlation = 0.82 ± 0.003; regression slope of 2.69, 95% CI [2.67, 2.71], intercept 1.8e-3, 95% CI [1.6e-3, 2.0e-3]). We observed similar performance gains when computing the eGRM using genealogies inferred by RELATE, with **EK**_relate_ attaining a highly correlated (r = 0.90 +/- 0.004; **Figure 2B**) and approximately unbiased estimate of **K**_all_ (regression slope of 0.96, intercept 3.7e-5; **Figure 2C**). We found **EK**_tsdate_ demonstrated lower correlation and biased estimates of **K**_all_ (**Figure 2B, C**). Taken together, our results suggest the eGRM is an unbiased and accurate estimator of the idealized canonical GRM containing all variants.

Next, we quantified the performance of eGRM when computed using genealogies inferred from a varying proportion of observed genetic variants. We found that the correlation between **K**_all_ and **EK**_relate_ was consistently higher than the correlation between **K**_all_ and **K**_obs_ (**Figure 2D** left). Moreover, for a fixed proportion of observed common SNPs (*e*.*g*., 20%; similar to SNP arrays), we observed the performance gap widened between **EK**_relate_ and **K**_obs_ as sample size increased (**Figure 2D** right). Intuitively, this improvement reflects the increasing contribution from rare variants to kinship in a larger sample that would not be captured by the canonical GRM based on only variants assayed on an array. Our results imply that eGRM can in principle more effectively capture relatedness in large-scale studies.

In practice, the construction of the GRM often uses imputed variants and/or is restricted to relatively common variants after pruning of correlated variants by LD. However, in simulations we found that pruning SNPs by LD before computing the GRM (**K**_obs (pruned)_) further decreased correlation with **K**_all_ (**Figure S2A**). When using imputed variants to construct the canonical GRM (**K**_obs (imputed)_), we found it more strongly correlated with **K**_all_ on average when compared with **K**_obs_ (**Figure S2A**). However, we observed performance in this scenario depends on relatedness between individuals in the imputation reference panel with target individuals, with correlation between **K**_obs (imputed)_ and **K**_all_ decreasing with average panel-relatedness (**Figure S2B**). The dependence on the availability of a closely related reference panel suggests that underrepresented populations would be at a disadvantage for genetic analysis using the canonical GRM^31^. Most importantly, across all of these scenarios, we observed our eGRM based on inferred genealogy (i.e., **EK**_relate_) consistently exhibited better correlation with **K**_all_ than **K**_obs (pruned)_ or **K**_obs (imputed)_ (**Figure S2A**).

### eGRM captures recent demographic events

Population structure, as the result of historical demographic processes that are encoded in the ARG, is conventionally visualized through principal components analysis (PCA) of the canonical GRM^4,32^. Because the eGRM is conditioned on the ARG encoding these historical events, we expected the eGRM to be more sensitive to population structure than the canonical GRM based on only common variants. To this end, we quantified the performance of the eGRM in capturing recent population structure through PCA. A. Motivated by recent work demonstrating that recent population structure (i.e. < 100 generations) is not well-captured by PCA computed from common SNP GRMs^29^, we simulated a stepping-stone population model with 25 demes spatially distributed in a 5×5 grid, with demes coalescing into a single deme 100 generations ago (**Figure 1B, Figure 3**; see **Methods**). We then compared the ability of the eGRM or the canonical GRM to identify recent population structure through PCA as quantified by the separation index (SI), which measures the proportion of neighbors in multi-dimensional space that are of the same deme or cluster (**Methods**).

**Figure 3.**
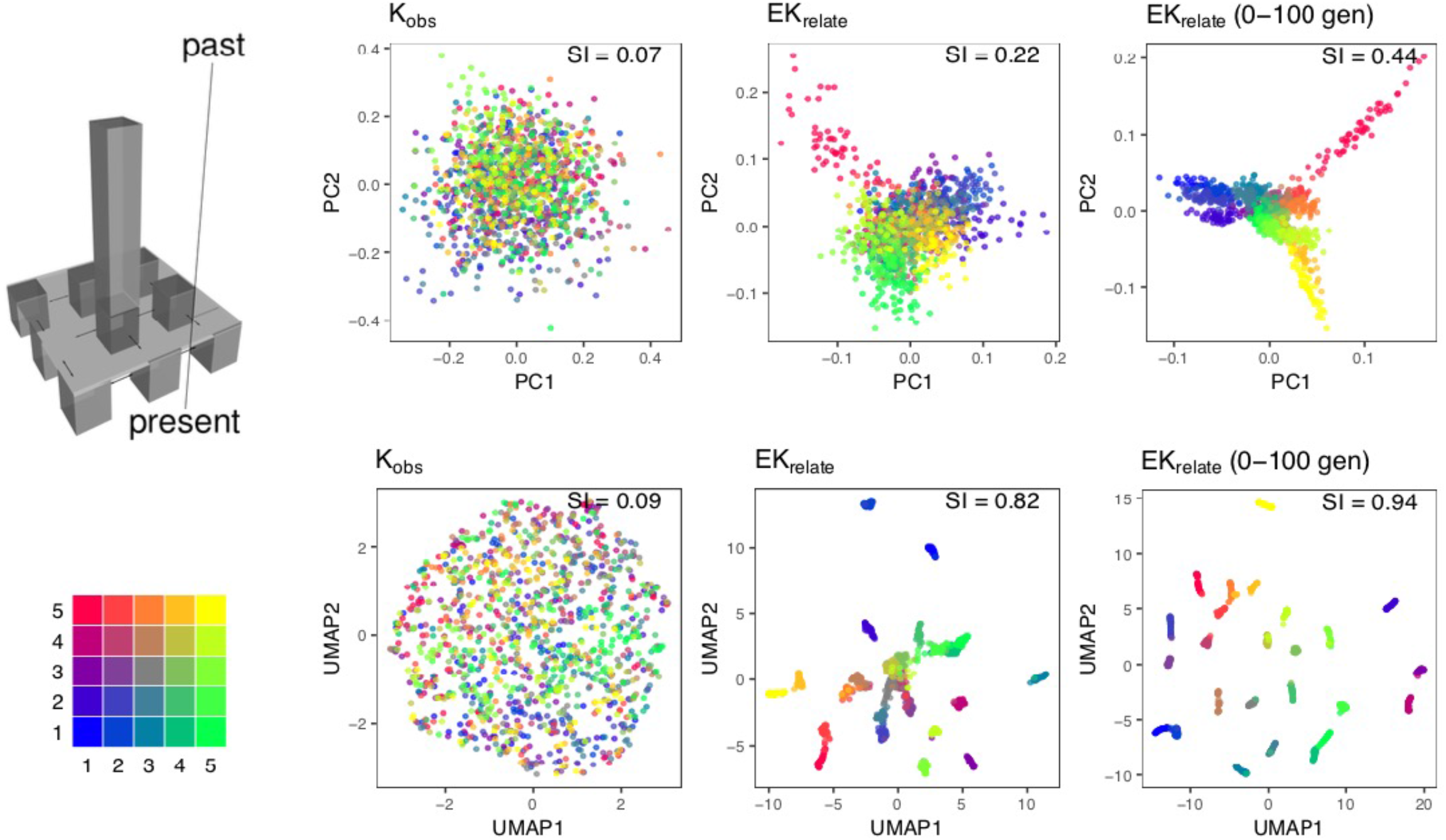
PCA based on eGRM more effectively captures recently established spatial structure in simulation compared to the canonical GRM. A 30Mb region was simulated, with 20% of common variants observed. Each deme has a constant population size of 500, in which 50 individuals are sampled. The first two PCs based on PCA of three GRMs are shown (top): the canonical GRM based on observed SNPs (**K**_obs_), the eGRM based on RELATE-reconstructed ARG using the observed SNPs (**EK**_relate_), or the partial eGRM based on the subset of branches between 0 and 100 generations across the ARG (**EK**_relate_ **(0-100 gen)**). Separation index (SI; see **Method** for a precise definition) is shown at the topright corner of each plot, which is the average proportion of same-label neighbors for each sample, indicating how well populations are separated. The first two features of UMAP transformation (bottom) applied to the top 10 PCs further accentuate the detected structure, as measured by SI.

To establish a baseline, we first recapitulated previous results demonstrating that PCA based on the GRM constructed using common variants (defined as MAF ≥ 0.05; **K**_common_), or the observed variants (defined as 20% of the variants, oversampling common variants; **K**_obs_), cannot distinguish the spatial structure of the demes (SI = 0.07-0.08 for **K**_common_ and **K**_obs_; **Figure 3, Figure S3A**). In comparison, the GRM constructed from rare variants (minor allele count = 2, 3, 4 or 5) alone (**K**_rare_; SI = 0.25) or all of the variants (**K**_all_; SI = 0.20; **Figure S3A**) can better detect structure.

We repeated this PCA analysis using **EK**_relate_ computed from the same set of variants used to construct **K**_**obs**_ and found it to better identify recent population structure (SI = 0.22; **Figure 3**). We next applied a UMAP transformation to the top 10 PCs based on each of the evaluated relatedness matrices, and found cluster separation performance improved when using **K**_rare_, **K**_all_, or **EK**_relate_ (SI = 0.80-0.82), but with little benefit when using **K**_common_ or **K**_obs_ (SI = 0.08-0.09; **Figure 3** and **Figure S3B**). We also simulated a demographic history with older population split times of 200 or 500 generations. In these scenarios, **EK**_relate_ (SI = 0.36 and 0.64 when split time is 200 and 500, respectively) consistently outperformed **K**_obs_ (SI = 0.13 and 0.34, respectively) in capturing population structure (**Figure S4**), likely due to additional haplotypic information captured by the inferred ARG, even though **K**_obs_ also improved in its performance capturing population structure than a scenario with more recent population split because the common variants are more informative of the population structure in these simulations. Therefore, under a structured model the eGRM consistently extract more information of the population structure compared to the canonical GRM based on the same set of variants.

### Partial eGRM reveals dynamic relatedness through history

By defining genome-wide relatedness as a function of coalescent trees, a major advantage of our eGRM framework is its natural generalization of relatedness constrained to a specific time window. We denote the eGRM computed from the ARG when considering only branches of a certain age as the *partial* eGRM (see **Methods**). To demonstrate the benefit of limiting relatedness calculation to certain generations, we re-analyzed our grid simulations but estimated **EK**_relate_ restricted to the most recent 100 generations (**Figure 3**). We observed that PCA based on **EK**_relate_ of the most recent 100 generations improved its ability to delineate population structure (SI = 0.44 for, compared to SI = 0.22 for **EK**_relate_ or 0.25 for **K**_rare_; **Figure 3, Figure S3A**). We found performance to further improve when applying an additional UMAP transformation on PCs (SI = 0.94; **Figure 3**).

The partial eGRM also provides new insight into the limitation of the canonical GRM method. In our single population simulation, GRM (**K**_**obs**_) was best correlated with very ancient partial eGRM, because the allelic ages of observed SNPs are generally much older (mean age of 12,872 generations; **Figure S5A, D**). In our grid structure simulations, due to smaller population sizes and stronger genetic drift, the common SNPs are much younger (mean age of 2,089 generations), resulting in **K**_obs_ having the highest correlation with partial eGRM between 2000-3000 generations ago (**Figure S5B, D**). The correlation with partial eGRM decreased going further back in time, likely due to older variants becoming fixed in the population and thus being excluded from **K**_obs_ computation. The expansion into multiple demes only occurred 100 generations ago, but because the allelic ages of observed SNPs used to construct **K**_obs_ generally predated this event (**Figure S5D**), individually these SNPs contained little information for the recent structure. Taken together, these results indicate that the canonical GRM provides a coarse measure of relatedness in which older, common SNPs are enriched, while the eGRM and partial eGRM provide a more fine-grained measure of relatedness at different time points.

### eGRM improves prediction of geographical pattern in empirical data

We evaluated the ability of eGRM to detect population structure in real world data by applying the framework to the genotyping array data of a Finnish cohort, FinMetSeq. We computed the canonical GRM and **EK**_relate_ based on 208,681 SNPs genotyped on 2644 individuals with both parents born in the same municipality in Finland^21^ (**Methods**). Using parental birthplaces as population labels, we found PCA of **EK**_relate_ was able to identify patterns of structure (SI = 0.52; **Figure 4A**) whereas the canonical GRM displayed mild separation between individuals with recent ancestry along differing regions of Finland (SI = 0.39; **Figure 4A**). Lower-order PCs computed from **EK**_relate_ revealed additional structure not matched by **K**_obs_ (SI = 0.47 vs SI = 0.29 for PC3-PC6; **Figure S6**). Notably, the first two PCs of **EK**_relate_ were explained in part by individuals with both parents born in the Surrendered Karelia (magenta color in **Figure 4**). Similar to our simulated results, when we applied UMAP to top PCs we observed improved resolution of fine-scale structure, with UMAP based on **EK**_relate_ continues to be more informative of the fine-scale structure within Finland than that based on the canonical GRM, regardless of the number of PCs included in UMAP (**Figure 4B**).

**Figure 4.**
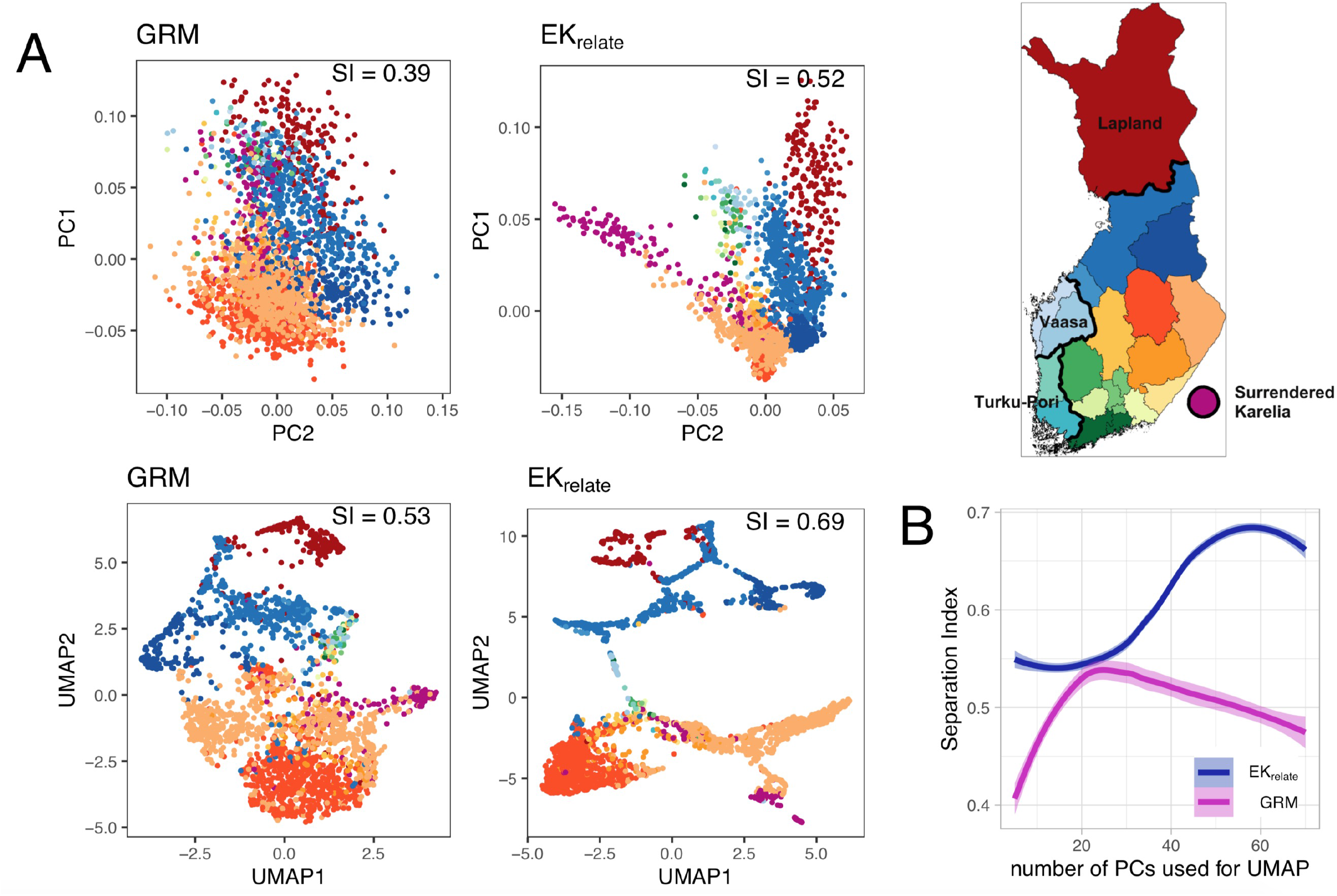
Clustering analysis based on eGRM revealed novel population structure in the population of Northern and Eastern Finland. (**A)** PCA and PCA+UMAP based on either **K**_obs_ or **EK**_relate_ are shown. A map of Finland with regions colored is provided for reference. Main geographical locations referenced in the text are labeled (Surrendered Karelia colored in magenta, Lapland colored in red, Turku-Pori colored in light green, and Vassa colored in light blue). Scatterplot of the first two features of UMAP transformation was based on the first 24 and 58 components of PCA of **K**_obs_ and **EK**_relate_, respectively. These numbers were chosen as they respectively are the number of components at which the separation index (SI) is maximized after applying the UMAP transformation. (**B**) Separation index achieved as successive PCs were included in UMAP transformation of PCA of **K**_obs_ and **EK**_relate_.

Next, to shed light on historical migration and population movements for FinMetSeq data, we computed and analyzed the partial eGRM considering only branches for the past 0-100 generations (**Figure S5C**). PCA of the partial eGRM suggests that recent structure in Northern and Eastern Finland is mainly driven by individuals from Lapland (colored red in **Figure 4**; the Northernmost part of Finland and home to the indigenous Finno-Urgic people, Sami), Surrendered Karelia (magenta), and Turku-Pori and Vaasa (light green and light blue, sharing major port borders with Sweden). Computing a partial eGRM further in the past exhibited patterns more similar to those found in the canonical GRM (**Figure S5C**). Qualitatively, the pattern of the partial eGRM at varying time depth and its correlation with a fixed canonical GRM are more reminiscent of the pattern observed in the grid structure simulation (**Figure S5B**) than the pattern in a single homogenous population (**Figure S5A**). Together these findings further support previous claims that common variants are enriched for those that survived a bottleneck in Finland and that there are extensive internal structure due to recent population movement, isolation, and drift^21,33–35^.

### Time and memory considerations for eGRM algorithm

We implemented the eGRM in a flexible Python framework using custom C extensions to accelerate core eGRM calculations. Our implementation of eGRM is memory-efficient. The main memory usage throughout the algorithm is a matrix of size *N* × *N*, which takes 8*N*^2^ bytes of memory when stored as doubles in C. However, outputting the resulting matrix into a NumPy array dominates the overall memory consumption (**Figure S7**). The time cost of computing eGRM is *Θ*(*mN*^2^), where *m* is the number of genealogical trees (see **Supplementary Text**). In the case of the FinMetSeq data, the genome-wide eGRM takes ∼30 hours on a single CPU to compute for 2644 samples and ∼120,000 genealogical trees.

## Discussion

In the current study we introduce the eGRM, a genealogical estimate of genetic relatedness. The eGRM is conceptually distinct from the canonical GRM, which is variant- or mutation-centric. As a result, analyses utilizing the canonical GRM need to be interpreted within the context of the marker ascertainment. Ascertainment could be biased because of availability of data, technical errors in data generation, or inconsistent analytical conventions across analysts. In contrast, the eGRM does not depend directly on the detection of variation (eGRM based on the true ARG does not depend on variation; but haplotypes based on a set of variant is used for inferring the ARG in practice), and thus is more robust when used in analyses with incomplete data.

A number of methods have been proposed to exploit the rich information stored in the ARG to make inference of population genetic parameters (*e*.*g*. for selection^36,37^ or population history^38,39^). Similarly, recent theoretical work has demonstrated the relationship between mutational processes by site, or on branches and nodes of the ARG^40^. Given the ARG, our framework considers mutations as appearing uniformly at random on the ARG, and relatedness between pairs of individuals is based on the probability of shared mutation, which is proportional to the branch lengths relating the two individuals. Our decision to explore this framework over alternative paths such as manipulating a matrix of TMRCAs is driven by the conceptual shift in treating mutations as random. We expect that our genealogical framework to compute genetic relatedness will enable seamless incorporation with downstream statistical applications, such as its inclusion in a linear mixed model for controlling population structure in association testing.

Through extensive simulations, we demonstrated that the eGRM is highly correlated with TMRCA and importantly provides an approximately unbiased estimate of **K**_all_. The former we used as the standard for true relatedness given a single tree, while the latter is an idealized GRM assuming all variants are perfectly observed. We illustrated the improvement and new insights that could be garnered using the eGRM with an application in the detection of population structure. First, common SNPs were thought to be uninformative about recent population structures because they tend to predate recent population divergence and are likely shared across all populations^29^. However, since haplotypes derived from common variants are used to infer the ARG, we showed that the eGRM detects the recent split and can better separate the spatial structure among demes, despite relying on only the common SNPs. Second, our framework provides a means to flexibly probe into the population structure at arbitrary time depths through the partial eGRM, suggesting that it can be optimized to account for structure at varying time scales. Third, we demonstrated empirically the insights of population structure that can be learned from eGRM using Finnish genotyping data. In contrast to the GRM, PCA and PCA+UMAP based on the eGRM could better delineate subpopulations such as individuals from the Surrendered Karelia region or from the southwestern region of Finland. The Surrendered Karelia region was a geographical region at the border of Finland and Russia but was ceded to Russia in 1940. Finnish citizens in this area were evacuated and resettled throughout the rest of Finland^41^. Contribution to the population structure of Finnish due to the evacuees and their descendants would not have been apparent if we examined only the PCA based on the canonical GRM (**Figure 4A**). Finally, by examining the partial eGRMs, the structure in Northern and Eastern Finland appeared to be more recently established, and the pattern of variation from this dataset is more consistent with a recently structured population with enhanced drift, rather than the conventional belief that Finnish composes a single homogeneous population^21,42^.

We have found that eGRM inference is stable when using computationally reconstructed genealogies rather than the true ARG (*e*.*g*., **EK**_relate_). However, we note that underlying assumptions required for accurate ARG inference may not be met in massive sample sizes. For example, it is often assumed that there are no recurrent mutations and multiple coalescent events per generation; both of these assumptions would be violated in extremely large samples. Mutation rate is often assumed to be constant across all branches of the inferred tree, which may not be true empirically^43^, and ARG inferences currently may not appropriately model all recombination events^44^. The extent to which relaxation or violation of these assumptions impact the ARG inference and its downstream computation of the eGRM will need to be evaluated systematically. Furthermore, a major current impediment is the scalability of ARG inference. In our simulations, eGRM from RELATE-reconstructed ARGs performs better than that of TSINFER+TSDATE, perhaps due to the inferred branch length information in RELATE, but the computational time is also orders of magnitude longer. As a result, even though the eGRM computation usually takes less than 5% of the total runtime, we were unable to efficiently compute eGRM on RELATE-reconstructed ARGs beyond 10,000 diploid individuals. Nevertheless, computational advances may well continue to make ARG inferences more accurate and scalable; until recently ARG inferences were restricted to only tens of individuals. As faster or more accurate ARG inference algorithms become available, our method will be primed to achieve advanced usability and performance with little adjustment.

Even without a more scalable ARG inference methods, eGRM will have the potential to make an immediate impact in genetic studies of humans and other species. For instance, most understudied populations are not resourced with a matched imputation reference panel or whole-genome sequencing data^31,45^. Even when genotyping array data are available, the arrays are rarely designed to represent variation found in the population of interest^46,47^. The biased ascertainment of incomplete genomic information is anticipated to exacerbate the disparity in our understanding of the genetic architecture between different populations. The eGRM could overcome these limitations because it is able to improve relatedness estimation, using only a subset of common markers nonetheless, to a level comparable to the canonical GRM constructed in presence of a population-matched imputation reference. The eGRM could thus enable analysis of limited genetic data and genetic mapping studies from under-resourced populations. Stepping outside of human studies, the genetic studies of other ecological species are rarely equipped with complete genomic information. In some cases, complete genomes of a sample are impossible to obtain, such as phylogenetic or ancestral studies on historical specimens. However, ARG could be inferred from limited genotyping data^48^, suggesting the eGRM can fill in the void in these studies.

## Methods

### Pairwise genetic relatedness with unobserved markers

We first describe the canonical and expected GRM (eGRM) in a haploid scenario; our framework can easily be generalized to diploid scenarios as describe in **Supplementary Methods**. We model the haplotypes of *N* samples with *M* variants as an *N* × *M* binary matrix *X* and denote the *N* × 1 vector at the *k*-th variant as *X*_*k*_. The sample allele frequency is 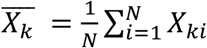 and it is required that 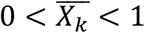 for *k* to be a variant site. Given *X*, the genetic relationship matrix (GRM) is commonly defined as

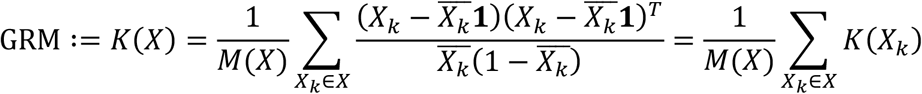

where **1** is the all-ones vector and *M*(*X*) is the number of variants in *X*. However, in practice only *M*′ < *M* markers are observed, resulting in the *N* × *M*′ observed haplotype matrix *X*′. The GRM computed using *X*′ is given by *K*(*X*′). As its definition suggests, *K*(*X*′) reflects the pair-wise relatedness conditioned on the observed haplotype data *X*′, and provides an incomplete picture of relatedness measure between individuals. Even though the complete haplotype matrix *X* is unknown (*M* is also unknown), through linkage between the unobserved markers in *X* and observed markers in *X*′, there may exist a more reasonable estimate of *K*(*X*) than *K*(*X*′) itself. Specifically, we show that when the ancestral recombination graph (ARG) *G* that connects the *N* samples is given, the expectation of *K*(*X*) can be derived, which we denote as the expected GRM (eGRM) on *G*.

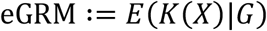

In practice, the ancestral recombination graph *G* can be inferred from the observed haplotypes *X*′ using recently emerging genealogical tree reconstruction tools^27,28^. Intuitively, we wish to define an eGRM where each entry represents the expected similarity between a pair of individuals should a mutation arise randomly in the ARG, after accounting for the expected similarity between a random pair of individuals in the same context given the structure of the ARG.

### Expectation of pairwise relatedness given a genealogy

The recombination and coalescent history of the *N* haploid samples can be completely represented by the ancestral recombination graph^25,26^ (ARG) denoted as *G*, which consists of a sequence of genealogical trees *G* = {*T*_*j*_|1 < *j* < *m*} across the whole genome. Each tree *T*_*j*_ = (*V*_*j*_, *E*_*j*_) is a directed binary tree with each node *v* ∈ *V*_*j*_ representing a chromosomal segment of a sample or an ancestor, and each branch *e* ∈ *E*_*j*_ representing the history of its child node until it coalesced into its parent node. We define *x*(*e*) as the haplotype vector associated with *e*, that is

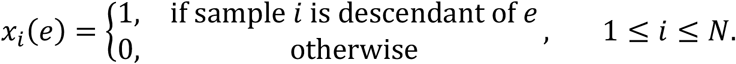

We assume *G* is fixed, and *X* is generated randomly through mutations occurring on *G*, implying expectation or variance over *X* is conditional on *G* by default. We overload set membership notation over *G* and denote that *e* is a branch in the ARG *G* as *e* ∈ *G*, to mean *e* ∈ *E*_*j*_ ∈ *T*_*j*_ ∈ *G* for some *j*.

Here, we define how mutations arise on *G*. For each branch *e* ∈ *E*_*j*_ we define *t*(*e*) as its length in generations, *l*(*e*) as the number of base pairs that *T*_*j*_ covers, and *u*(*e*) as the mutation rate on this branch. We model the number of mutations occurring on branch *e* as being Poisson distributed *M*(*e*)∼Pois(*μ*(*e*)) with rate *μ*(*e*) = *t*(*e*)*l*(*e*)*u*(*e*) which implies the total number of mutations over *G* also follows a Poisson distribution *M*∼Pois(*μ*(*G*)) with rate *μ*(*G*) = ∑_*e*∈*G*_ *μ*(*e*).

Next, we consider the sampling distribution of the complete haplotype matrix *X* and *K*(*X*) given *G*. All column vectors in *X* should be from {*x*(*e*)|*e* ∈ *G*}, and *x*(*e*) should have repeated *M*(*e*) times in *X*. We have

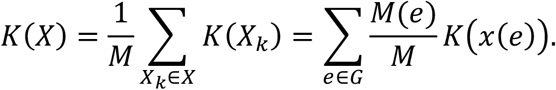

Note that *K*(*X*) can only be defined when *M* > 0. The expectation of *K*(*X*) is

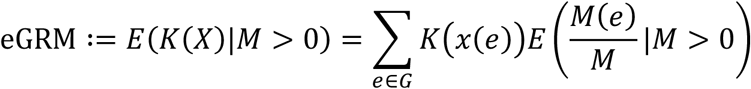

Using the fact that whenever *M* > 0, 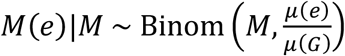, we can compute the expectation,

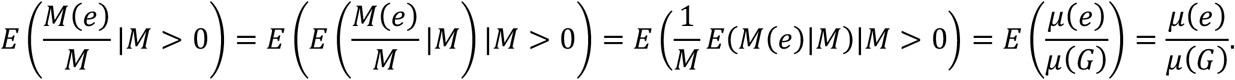

We have

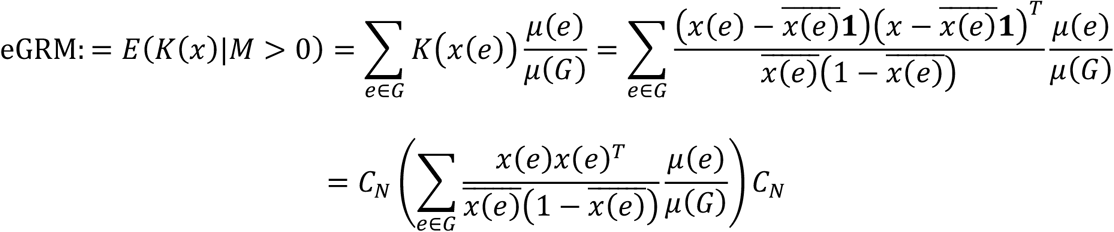

where 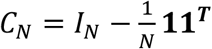 is a centering matrix, and *I*_*N*_ is the *N* × *N* identity matrix.

Computationally, we can compute eGRM by traversing *e* ∈ *G* in any order while updating a buffer matrix. For each branch *e*, we first compute 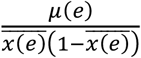 and add this number to a square submatrix of elements indexed by nonzero elements of *x*(*e*)*x*(*e*)^*T*^. Finally, we divide the resulting matrix by *μ*(*G*) and then center it by column and by row to get the eGRM of *G*.

Extension of the haploid eGRM to diploid organisms are straight-forward by considering and weighing the paternal and maternal haplotypes separately; details are provided in the **Supplementary Methods**. Furthermore, given our probabilistic formulation of relatedness given a genealogy, it is natural to define higher central moments. Thus, we also defined the element-wise Var(*K*(*X*)|*G*), which we term varGRM and captures the expected deviation around the individual entries in the eGRM, and explored its behavior briefly in **Figure S8**. Derivation of the varGRM can be found in **Supplementary Methods**.

### Genealogy and genotype simulations

We used two different demographic models in our simulation experiments (**Figure 1B**) for a comprehensive comparison between GRM and eGRM. To investigate the accuracy of eGRM compared with the canonical GRM in estimating true relatedness, we simulated genealogies and genotypes under a single-population exponential-growth model based on the published out-of-Africa demography^49^. Model parameters were suggested by the msprime documentation, based on the European branch of the model. We did not simulate the other two branches of population, nor the migration rates between the populations. To investigate the performance of eGRM compared with the canonical GRM in detecting recent population structure, we simulated genealogies and genotypes of a structured population with a 5×5 grid stepping-stone demographic model motivated by a similar model recently published by Zaidi and Mathieson^29^. We simulated 50 individuals per deme, with population size of 500 and migration rate of 0.01 per generation between neighboring demes. The 25 demes split from a single ancestral population of the same population size 100 generations ago.

We simulate genotypes and tree sequences by MSPRIME^50^, with mutation rate and recombination rates set to 1e-8 per bp per generation. To mimick observed genetic data derived from a genotyping array that biases towards the common variants, we restricted the observed set of variation to a subset (20% by default, unless otherwise specified) of the simulated variants with minor allele count ≥ 5. To oversample the common variants, we sampled with probability proportional to 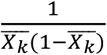, where 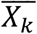, is the sample allele frequency of variant *k*. To show the practical use of eGRM, we reconstruct the sequence of trees from observed variants using RELATE^28^ and TSINFER+TSDATE^27,30^, using default parameters as suggested by the user manuals. The tree sequence output of RELATE is converted to TSKIT format, which contains a gap-filler tree with no genetic information between basepair zero and the first genetic marker in the dataset. In order to prevent overrepresenting a tree that covers a long region but with little actual information, we always skip the first tree in the tree sequence in our empirical analysis. We denote the canonical GRM based on observed variants as **K**_obs_, the eGRM computed using true genealogies as **EK**, and the eGRM computed using genealogies inferred from observed variants as input for RELATE or TSINFER+TSDATE as **EK**_relate_ or **EK**_tsdate_. Unless otherwise noted, by default all simulations are performed on a 30Mb chromosome with both mutation rates and recombination rates set to 10^57^ per generation per base pair.

### FinMetSeq genotyping and quality control

To exam the performance of eGRM on real genotyping data, we applied our method to a subset of the FinMetSeq dataset^21^ (dbGaP accession numbers: phs000743.v1.p1.c999, phs000756.v1.p1.c999) consisted of 2,644 samples who have self-reported that both parents were born in the same municipality in Finland. The dataset contained 1,504,461 SNPs from whole exome sequencing and a genotyping array backbone^21^. We retained only biallelic SNPs. We filtered variants with MAF ≥ 0.01 and missingness ≤ 0.01, resulting in 208,681 common SNPs. We phased the genotypes by EAGLE using its default hg19 genetic map. We reconstructed the genealogical tree sequence using RELATE with all parameters same as in its official manual. We then applied our eGRM algorithm on the resulting tree sequence to compute **EK**_relate_. The Canonical GRM was computed based on the same set of 208,681 SNPs after phasing with EAGLE.

### Population structure analysis

We contrasted and visualized the information of population structure contained in GRM and eGRM through principal components analysis (PCA) and uniform manifold approximation and projection (UMAP). PCA was computed using the “linalg.eig” function in the python “numpy” library, UMAP was computed by the R “umap” package with all default parameters. To quantitatively assess the improvement of eGRM over GRM in informing clustering analysis from structured populations, we devised a separation index to assess proportion of nearest neighbors that are in the same population in multi-dimensional space. Suppose we have a set of sample points *A* in a metric space with metric *d*. Each point *a* ∈ *A* has a true label *l*(*a*). The separation index defines *S*(*a*) = {*b* ∈ *A*|*l*(*b*) = *l*(*a*)} as the true class that *a* belongs. In simulated data, the true label is the deme or population membership of each individual. In empirical data, the birth place of the parents or grandparents was assumed to be the true label. We also define the size-*n* neighbor of *a* as the *n* nearest point of *a* including *a* itself, denoted as *R*_*n*_(*a*) = {*b* ∈ *A* such that |{*c* ∈ *A*|*d*(*b, a*) ≥ *d*(*c, a*)}| ≤ *n*}. The separation index is defined as the average proportion of same-class neighbors

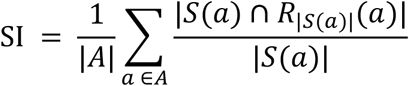

which is a real number between 0 and 1 indicating how well the metric *d* is capturing the true classification *l*. Note that SI is only dependent on the relative order of distances between pairs of points, making it a unified measure of clustering performance among PCA, UMAP and other distance-based methods.

### eGRM Software

We have implemented the algorithms related to eGRM in a python package ‘egrm’, which is publicly available in PyPI. Documentation of this package as well as simulation commands used in this study can be found on its github page (https://github.com/Ephraim-usc/egrm).

## Acknowledgement

We would like to thank Michael D. Edge, Diego Ortega-Del Vecchyo, Christian D. Huber, attendees of the 2020 Biology of Genomes, 2020 American Society of Human Genetics, and 2021 Probgen virtual meetings, and members of FinMetSeq consortium for discussions and help with data access. Research reported in this publication was supported by National Institute of General Medical Sciences (NIGMS) of the National Institute of Health under award number R35GM142783 (to C.W.K.C.). Computation for this work is supported by USC’s Center for Advanced Research Computing (https://carc.usc.edu).

## Author’s Contributions

C.W.K.C. conceived of the study. C.F., N.A.M. and C.W.K.C. designed the study. C.F. performed the analysis. C.F., N.A.M., and C.W.K.C. interpreted the data. C.W.K.C. contributed to the data collection. C.F., N.A.M., C.W.K.C. wrote the paper.

## Competing Interests

The authors declare no competing interests.

## Notes

### Competing Interest Statement

The authors have declared no competing interest.

## References

1. Chiang, C. W. K., Mangul, S., Robles, C. & Sankararaman, S. A Comprehensive Map of Genetic Variation in the World’s Largest Ethnic Group-Han Chinese. Mol Biol Evol 35, 2736–2750 (2018).

2. Novembre, J. et al. Genes mirror geography within Europe. Nature 456, 98–101 (2008).

3. Hirschhorn, J. N. & Daly, M. J. Genome-wide association studies for common diseases and complex traits. Nat Rev Genet 6, 95–108 (2005).

4. Patterson, N., Price, A. L. & Reich, D. Population structure and eigenanalysis. PLoS Genet 2, e190 (2006).

5. Price, A. L. et al. Principal components analysis corrects for stratification in genome-wide association studies. Nat Genet 38, 904–9 (2006).

6. Kang, H. M. et al. Variance component model to account for sample structure in genome-wide association studies. Nat Genet 42, 348–54 (2010).

7. Lippert, C. et al. FaST linear mixed models for genome-wide association studies. Nat Methods 8, 833–5 (2011).

8. Listgarten, J. et al. Improved linear mixed models for genome-wide association studies. Nat Methods 9, 525–6 (2012).

9. Loh, P. R. et al. Efficient Bayesian mixed-model analysis increases association power in large cohorts. Nat Genet 47, 284–90 (2015).

10. Zhou, X. & Stephens, M. Genome-wide efficient mixed-model analysis for association studies. Nat Genet 44, 821–4 (2012).

11. Yang, J. et al. Common SNPs explain a large proportion of the heritability for human height. Nat Genet 42, 565–9 (2010).

12. Speed, D. & Balding, D. J. Relatedness in the post-genomic era: is it still useful? Nat Rev Genet 16, 33–44 (2015).

13. Thompson, E. A. Identity by descent: variation in meiosis, across genomes, and in populations. Genetics 194, 301–26 (2013).

14. Powell, J. E., Visscher, P. M. & Goddard, M. E. Reconciling the analysis of IBD and IBS in complex trait studies. Nat Rev Genet 11, 800–805 (2010).

15. Visscher, P. M. et al. Assumption-Free Estimation of Heritability from Genome-Wide Identity-by-Descent Sharing between Full Siblings. PLoS Genet 2, e41 (2006).

16. Speed, D., Hemani, G., Johnson, M. R. & Balding, D. J. Improved Heritability Estimation from Genome-wide SNPs. The American Journal of Human Genetics 91, 1011–1021 (2012).

17. Mancuso, N. et al. The contribution of rare variation to prostate cancer heritability. Nat Genet 48, 30–5 (2016).

18. Hartman, K. A., Rashkin, S. R., Witte, J. S. & Hernandez, R. D. Imputed Genomic Data Reveals a Moderate Effect of Low Frequency Variants to the Heritability of Complex Human Traits. bioRxiv 2019.12.18.879916 (2019) doi:10.1101/2019.12.18.879916.

19. Hernandez, R. D. et al. Ultrarare variants drive substantial cis heritability of human gene expression. Nat Genet 51, 1349–1355 (2019).

20. Powell, J. E., Visscher, P. M. & Goddard, M. E. Reconciling the analysis of IBD and IBS in complex trait studies. Nat Rev Genet 11, 800–5 (2010).

21. Locke, A. E. et al. Exome sequencing of Finnish isolates enhances rare-variant association power. Nature 572, 323–328 (2019).

22. Chiang, C. W., Ralph, P. & Novembre, J. Conflation of Short Identity-by-Descent Segments Bias Their Inferred Length Distribution. G3 (Bethesda) 6, 1287–96 (2016).

23. Gusev, A. et al. Whole population, genome-wide mapping of hidden relatedness. Genome Res 19, 318–26 (2009).

24. Naseri, A., Liu, X., Tang, K., Zhang, S. & Zhi, D. RaPID: ultra-fast, powerful, and accurate detection of segments identical by descent (IBD) in biobank-scale cohorts. Genome Biol 20, 143 (2019).

25. Hudson, R. R. Gene genealogies and the coalescent process. (Oxford surveys in evolutionary biology, 1990).

26. Griffiths, R. C. & Marjoram, P. Ancestral Inference from Samples of DNA Sequences with Recombination. Journal of Computational Biology 3, 479–502 (1996).

27. Kelleher, J. et al. Inferring whole-genome histories in large population datasets. Nat Genet 51, 1330–1338 (2019).

28. Speidel, L., Forest, M., Shi, S. & Myers, S. R. A method for genome-wide genealogy estimation for thousands of samples. Nat Genet 51, 1321–1329 (2019).

29. Zaidi, A. A. & Mathieson, I. Demographic history mediates the effect of stratification on polygenic scores. eLife 9, e61548 (2020).

30. Wohns, A. W. et al. A unified genealogy of modern and ancient genomes. http://biorxiv.org/lookup/doi/10.1101/2021.02.16.431497 (2021) doi:10.1101/2021.02.16.431497.

31. Lin, M. et al. Population-specific reference panels are crucial for genetic analyses: an example of the CREBRF locus in Native Hawaiians. Hum. Mol. Genet. 29, 2275–2284 (2020).

32. McVean, G. A Genealogical Interpretation of Principal Components Analysis. PLoS Genet 5, e1000686 (2009).

33. Wang, S. R. et al. Simulation of Finnish population history, guided by empirical genetic data, to assess power of rare-variant tests in Finland. Am J Hum Genet 94, 710–20 (2014).

34. Martin, A. R. et al. Haplotype Sharing Provides Insights into Fine-Scale Population History and Disease in Finland. Am J Hum Genet 102, 760–775 (2018).

35. Kerminen, S. et al. Fine-Scale Genetic Structure in Finland. G3 (Bethesda) 7, 3459–3468 (2017).

36. Stern, A. J., Wilton, P. R. & Nielsen, R. An approximate full-likelihood method for inferring selection and allele frequency trajectories from DNA sequence data. PLoS Genet 15, e1008384 (2019).

37. Stern, A. J., Speidel, L., Zaitlen, N. A. & Nielsen, R. Disentangling selection on genetically correlated polygenic traits via whole-genome genealogies. Am J Hum Genet (2021) doi:10.1016/j.ajhg.2020.12.005.

38. Li, H. & Durbin, R. Inference of human population history from individual whole-genome sequences. Nature 475, 493–6 (2011).

39. Schiffels, S. & Durbin, R. Inferring human population size and separation history from multiple genome sequences. Nat Genet 46, 919–25 (2014).

40. Ralph, P., Thornton, K. & Kelleher, J. Efficiently Summarizing Relationships in Large Samples: A General Duality Between Statistics of Genealogies and Genomes. Genetics 215, 779–797 (2020).

41. Armstrong, K. Remembering Karelia: a family’s story of displacement during and after the Finnish wars. (Berghahn Books, 2004).

42. Jakkula, E. et al. The genome-wide patterns of variation expose significant substructure in a founder population. Am J Hum Genet 83, 787–94 (2008).

43. Harris, K. & Pritchard, J. K. Rapid evolution of the human mutation spectrum. eLife 6, e24284 (2017).

44. Deng, Y., Song, Y. S. & Nielsen, R. The distribution of waiting distances in ancestral recombination graphs. Theoretical Population Biology S0040580921000484 (2021) doi:10.1016/j.tpb.2021.06.003.

45. Xu, Z. M. et al. Using population-specific add-on polymorphisms to improve genotype imputation in underrepresented populations. http://biorxiv.org/lookup/doi/10.1101/2021.02.03.429542 (2021) doi:10.1101/2021.02.03.429542.

46. Martin, A. R. et al. Low-coverage sequencing cost-effectively detects known and novel variation in underrepresented populations. http://biorxiv.org/lookup/doi/10.1101/2020.04.27.064832 (2020) doi:10.1101/2020.04.27.064832.

47. Wojcik, G. L. et al. Imputation-Aware Tag SNP Selection To Improve Power for Large-Scale, Multi-ethnic Association Studies. G3 (Bethesda) (2018) doi:10.1534/g3.118.200502.

48. Speidel, L. et al. Inferring population histories for ancient genomes using genome-wide genealogies. http://biorxiv.org/lookup/doi/10.1101/2021.02.17.431573 (2021) doi:10.1101/2021.02.17.431573.

49. Gutenkunst, R. N., Hernandez, R. D., Williamson, S. H. & Bustamante, C. D. Inferring the Joint Demographic History of Multiple Populations from Multidimensional SNP Frequency Data. PLoS Genet 5, e1000695 (2009).

50. Kelleher, J., Etheridge, A. M. & McVean, G. Efficient Coalescent Simulation and Genealogical Analysis for Large Sample Sizes. PLoS Comput Biol 12, e1004842 (2016).

