## Supplemental Figures for "A genealogical estimate of genetic relationships"

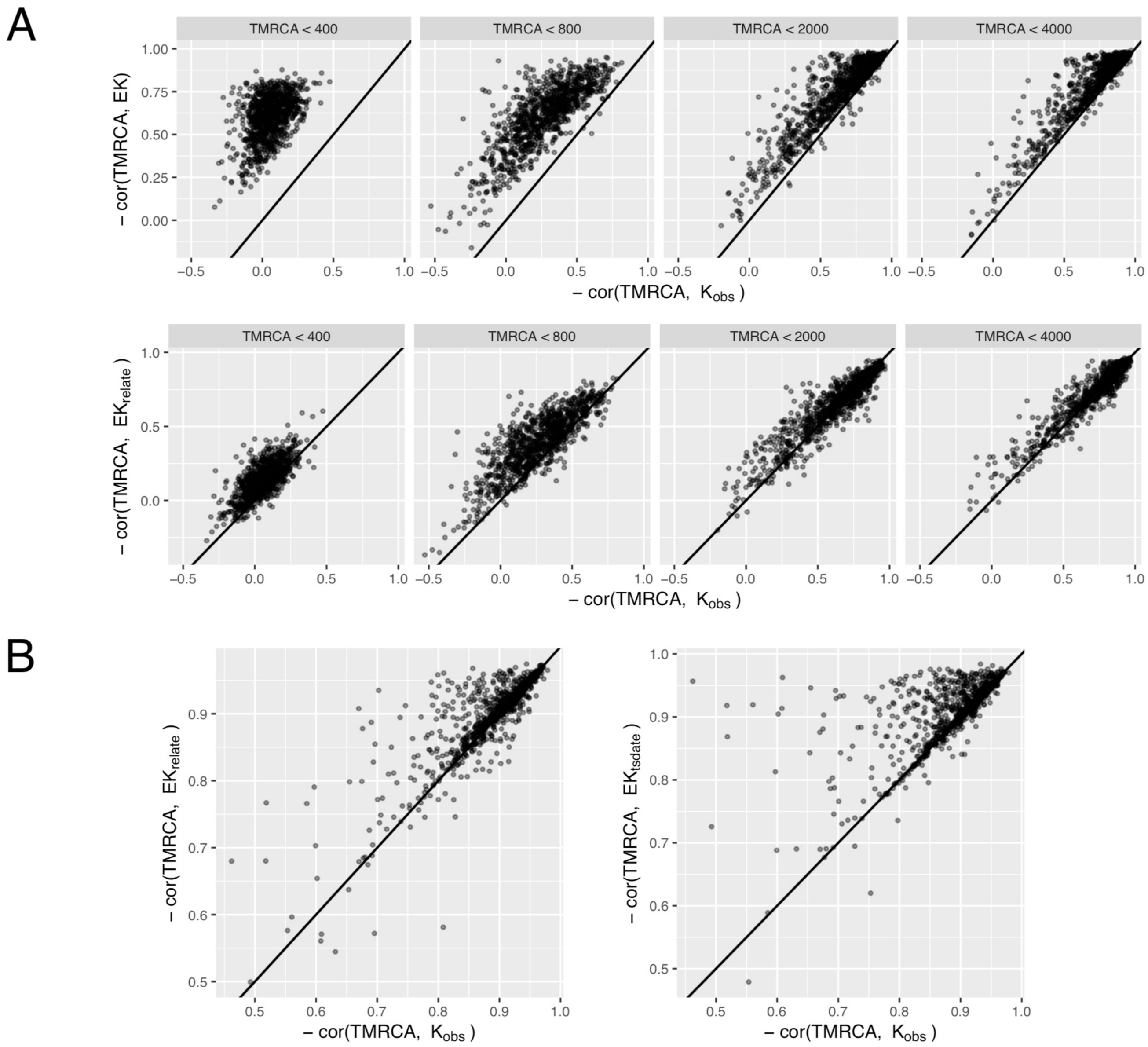


*Figure S1*

**eGRM shows stronger correlation with TMRCA than the canonical GRM across multiple time slices.** (**A**) Negative Spearman correlation between TMRCA and **EK** (top row) or $\mathbf{EK_{relate}}$ (bottom row), restricted to pairs of individuals with TMRCA less than the cutoff values, as compared to the correlation between TMRCA and $\mathbf{K_{obs}}$. In both cases, **EK** or $\mathbf{EK_{relate}}$ are better correlated with TMRCA than $\mathbf{K_{obs}}$ at more recent time scale. (**B**) Negative Spearman correlation between TMRCA and $\mathbf{EK}_{relate}$ (left) and $\mathbf{EK}_{tsdate}$ (right). In all cases the correlation between TMRCA and each eGRM is better than that between TMRCA and $\mathbf{K}_{obs}$. The correlation is better between TMRCA and eGRM 73.6% (P = 4e-52) and 76.5% (P = 5e-66) of the time, for $\mathbf{EK}_{relate}$ and $\mathbf{EK}_{tsdate}$, respectively.

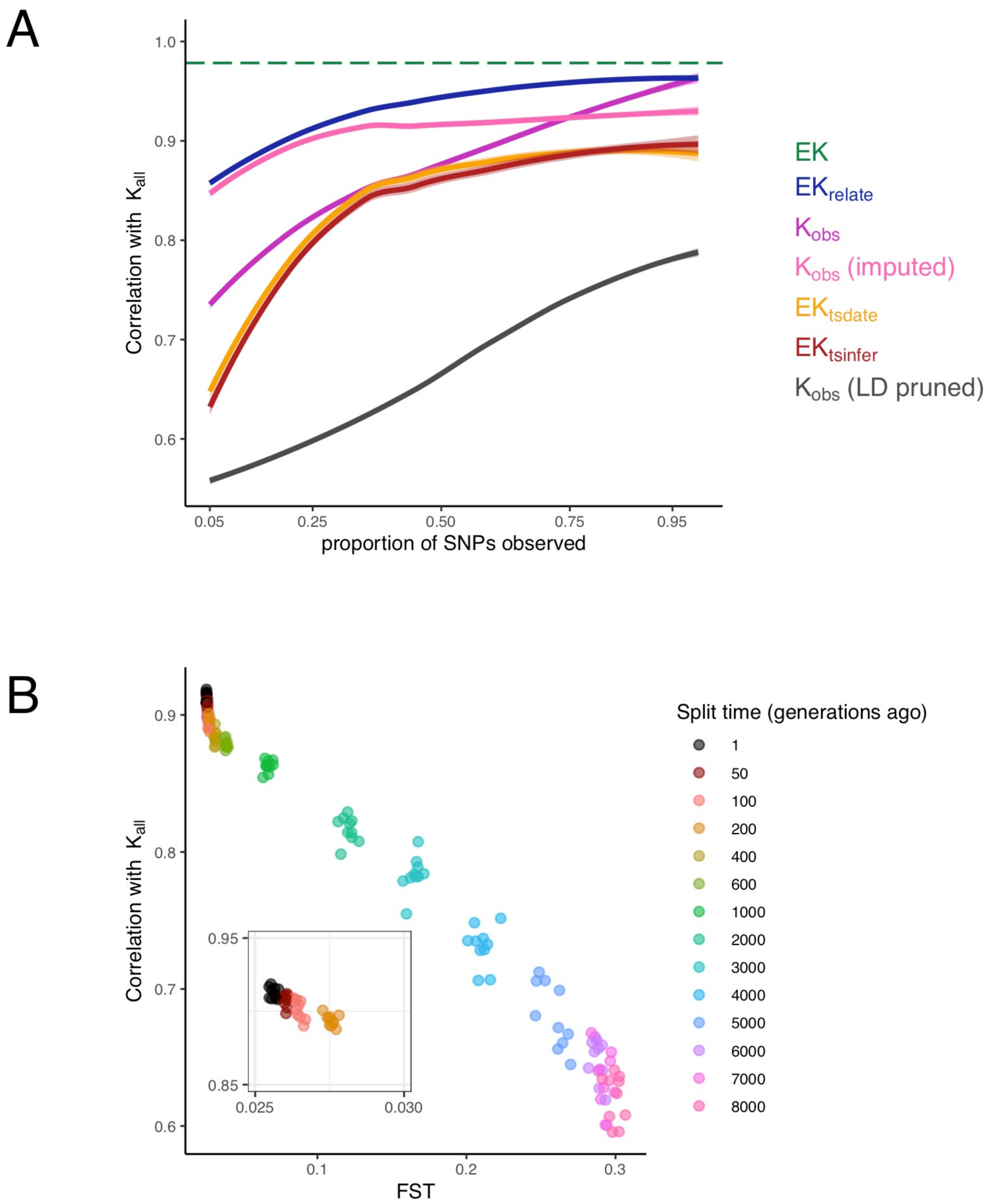


*Figure S2*

**Effectiveness of the canonical GRM based on imputed or pruned variants to capture relatedness.** (**A**) Pearson correlation with $\mathbf{K}_{all}$ with varying proportion of SNPs observed. $\mathbf{K}_{obs}$ **(imputed)** denotes the GRM based on observed variants (20% of all variants over-sampling the common variants, **Methods**) after imputation using reference samples (N = 1000) from a separate population diverged 100 generations ago (resulting in an $F_{ST}$ = 0.027 relative to the study population). $\mathbf{K}_{obs}$ (LD pruned) denotes GRM based on observed variants after pruning the set of observed variants by PLINK2.0 using the parameter "--indep-pairwise 50 5 0.1". $\mathbf{EK}_{tsdate}$ denotes eGRM based on TSINFER+TSDATE reconstructed ARG; TSDATE was run with effective population size 10,000, mutation rate 1e-8, and otherwise default parameters. The remaining matrices included in the comparison are the same as that shown in **Figure 2**. (**B**) Pearson correlation between $\mathbf{K}_{all}$ and imputed $\mathbf{K}_{obs}$ using reference panels from a separate population with varying divergence times. The horizontal axis shows the $F_{ST}$ between the reference and the study populations computed based on 20 randomly sampled individuals from each population. For reference, $F_{ST}$ between human non-African populations within the same continent are generally < 0.05; $F_{ST}$ between African populations or between populations across continents are generally < 0.2 (ref. [43]).

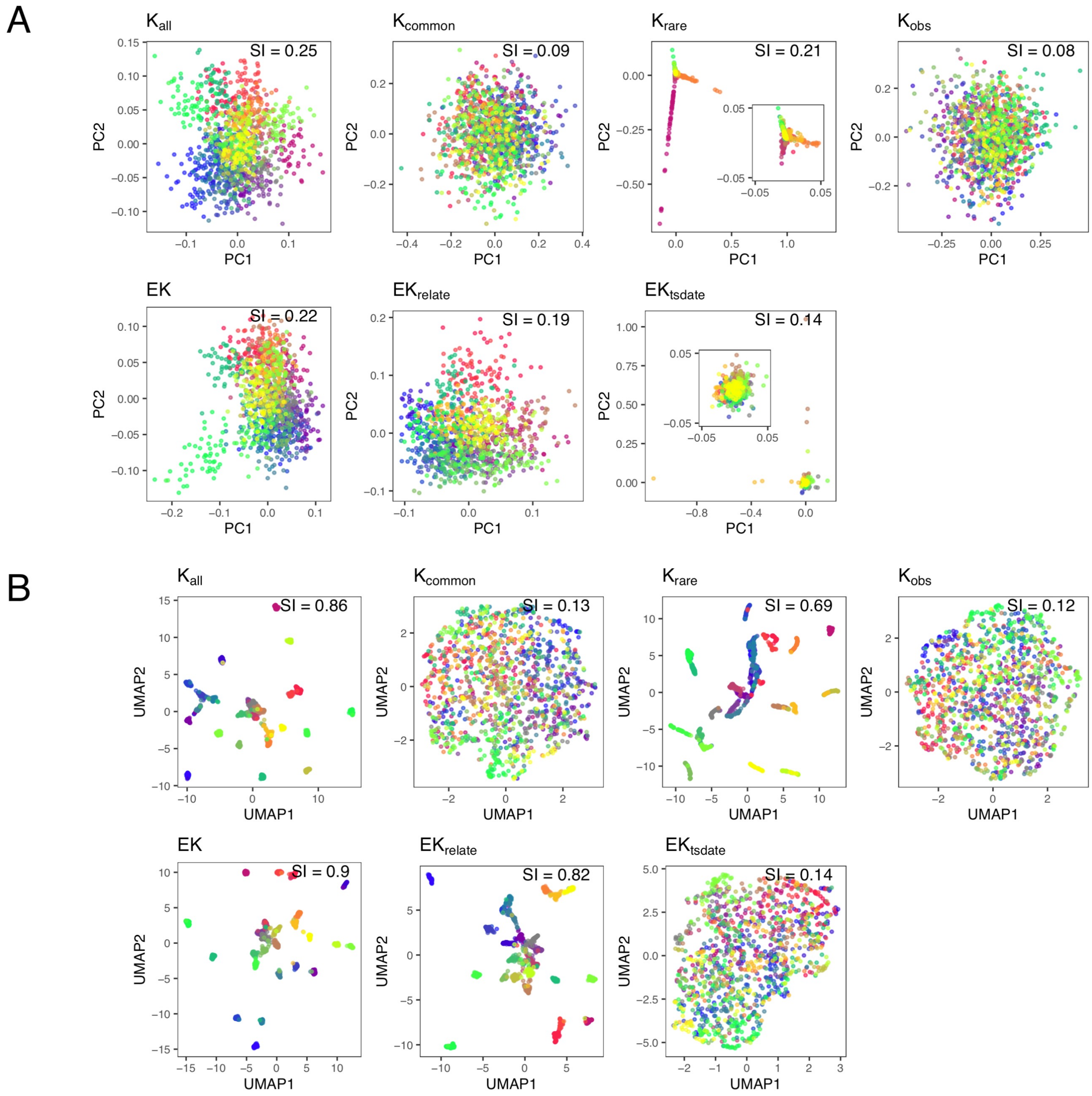


*Figure S3*

**Performance of various GRM and eGRM in detecting population structure in the grid-like recent expansion model.** (**A)** First two components of PCA on the canonical GRM and **EK**$_{relate}$. **K**$_{common}$ is computed based on all variants in simulation with MAF ≥ 0.05; **K**$_{rare}$ is based on all variants with minor allele count = 2,3,4 or 5. Insets provide a zoomed in view of the largest mass of datapoints in the PCA. (**B**) First two features of the UMAP transformation applied to the top 10 PCs from **A**. Simulations were done in the same manner as **Figure 3**.

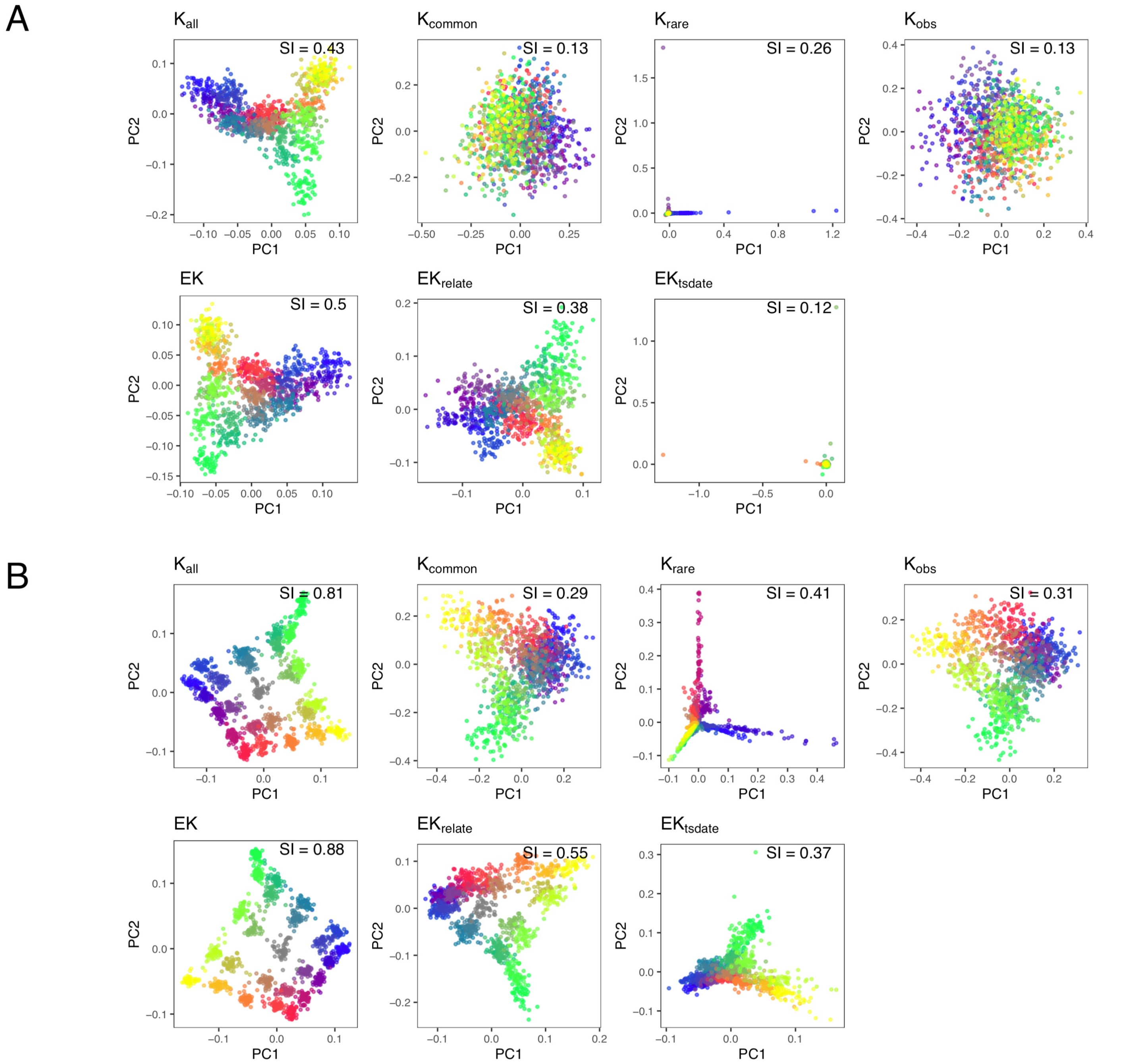


*Figure S4*

**Performance of various GRM and eGRM in detecting population structure in the grid-like model with varying divergence times.** The first two components of PCA on the indicated GRM or eGRM in the grid-like simulation where the divergence time between demes were set at (**A**) 200 or (**B**) 500 generations ago. In (**A**), **$K_{obs}$** began to be informative of population structure while **$EK_{relate}$** was able to capture the correct overall structure of the four corners of the 5x5 spatial grid. In (**B**), **$K_{obs}$** also began to capture the overall structure while **$EK_{relate}$** was able to identify each deme. In contrast, **$K_{rare}$** appears to consistently provide only information of the most recent past, and in (**B**) would be less effective than **$EK_{relate}$** to detect the structure as measured by SI.

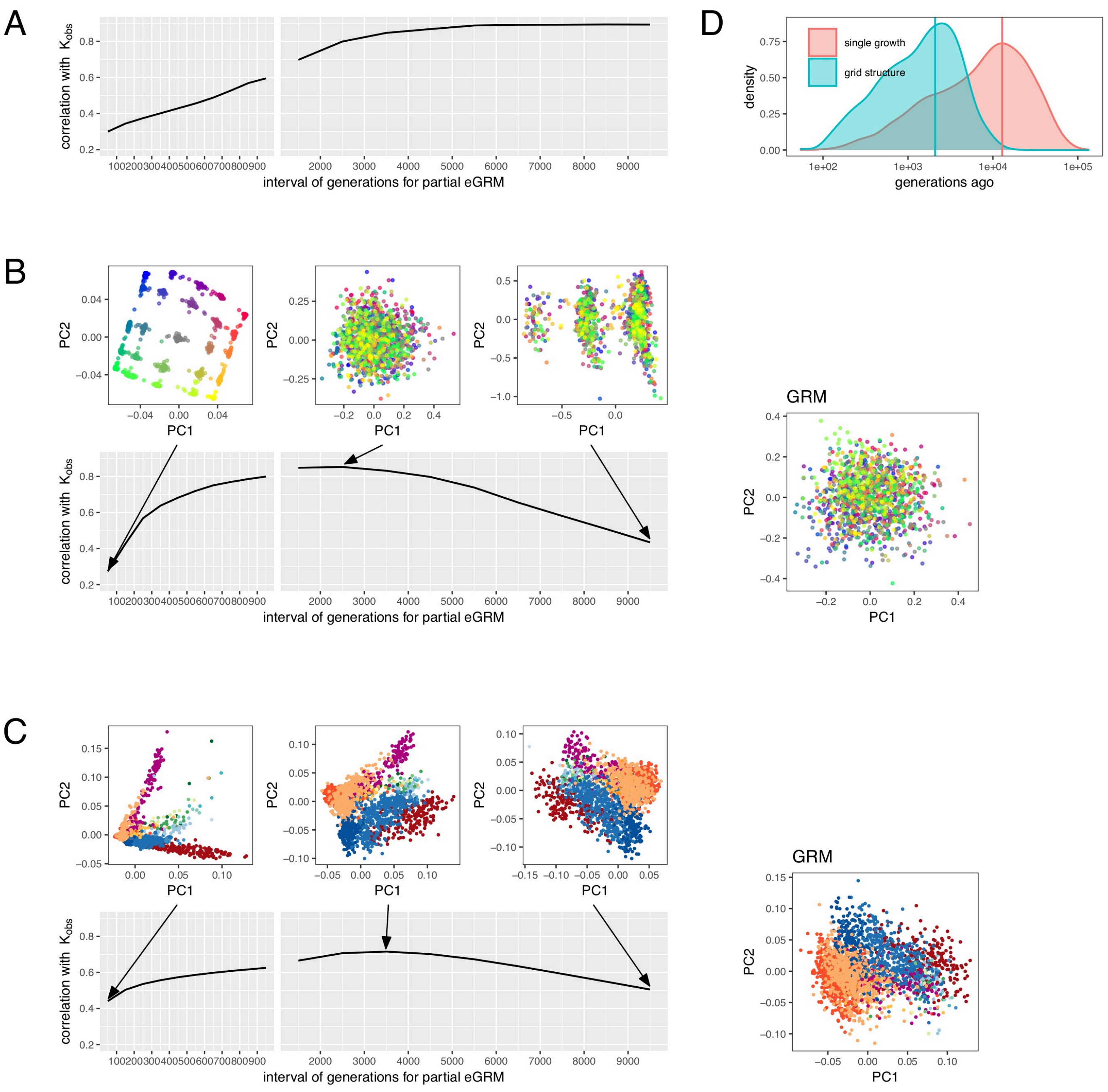


*Figure S5*

**Partial eGRM profile changes in population structure over time.** Correlation between $\mathbf{K_{obs}}$ and partial eGRM between different time intervals in (**A**) the single-growth simulations, (**B**) the grid structure simulations, and (**C**) the FinMetSeq data. Partial eGRM were constructed in intervals of 100 generations up to generation 1,000, and in intervals of 1000 generations up to generation 10,000. In (**B**), PCA plots of partial **EK** between 0-100, 2000-3000 and 9000-10000 generations are shown on top, which illustrates the clearly detectable changes in population structure over time if the true ARG were known. In (**C**), PCA plots of partial $\mathbf{EK}_{relate}$ between 0-100, 3000-4000 and 9000-10000 generations are shown on top. This profile of correlation of $\mathbf{K_{obs}}$ and the empirial partial eGRM from the FinMetSeq data is qualitatively more similar to the grid structure simulation in (**B**) rather than a single growth simulation in (**A**). (**D**) Distribution of allele age in generations of observed variants in the single-growth and grid structure simulations. In the grid structure model, observed variants are from 2089 generations on average, which explains that $\mathbf{K}_{obs}$ correlates best with 2000-3000 generations partial **EK**. In the single-growth model, however, observed variants are much older due to a larger population size, so that $\mathbf{K}_{obs}$ correlates better with very ancient partial **EK**.

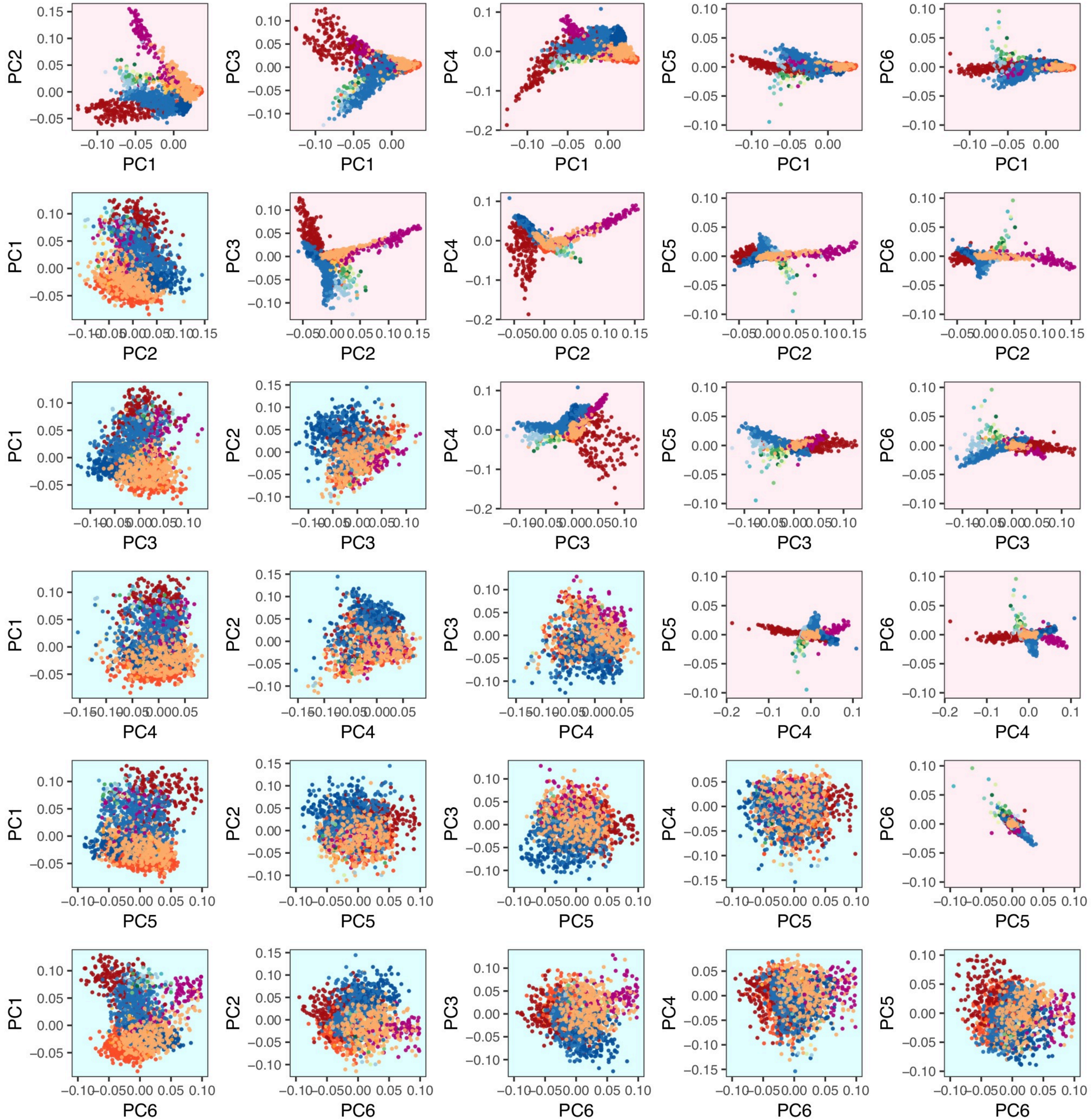


*Figure S6*

**Pairwise comparisons of PCs 1-6 based on $\mathbf{K}_{obs}$ or $\mathbf{EK}_{relate}$ from the FinMetSeq data**. All pairwise combinations of bivariate PC plots for PCs 1 through 6 are shown for $\mathbf{K}_{obs}$ in the lower triangle, and for $\mathbf{EK}_{relate}$ in the upper triangle. In general, additional structure in the FinMetSeq population is still being revealed by lower PCs of $\mathbf{EK}_{relate}$ as samples from different geographical locations continue to differentiate among lower PCs. In contrast, By PC4 of $\mathbf{K}_{obs}$ there is no longer discernable geographical structure.

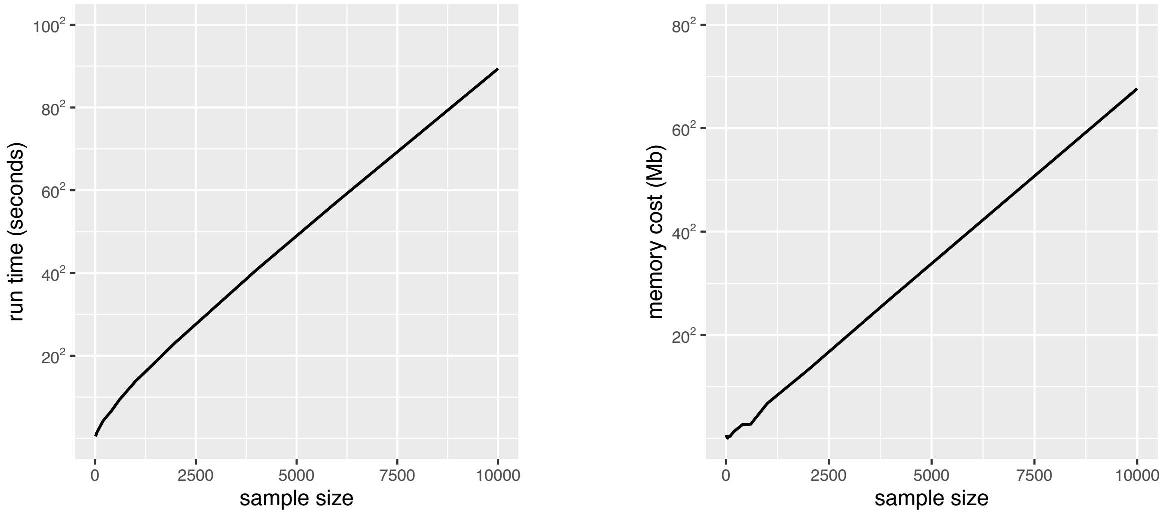


*Figure S7*

**Benchmarking the computation time and memory cost of eGRM algorithm on the true ARG of a 30Mb chromosome.** The run time in seconds and memory consumption in Mb are shown for computing the eGRM based on the true ARG from simulation of 1000 individuals from the single growth model.

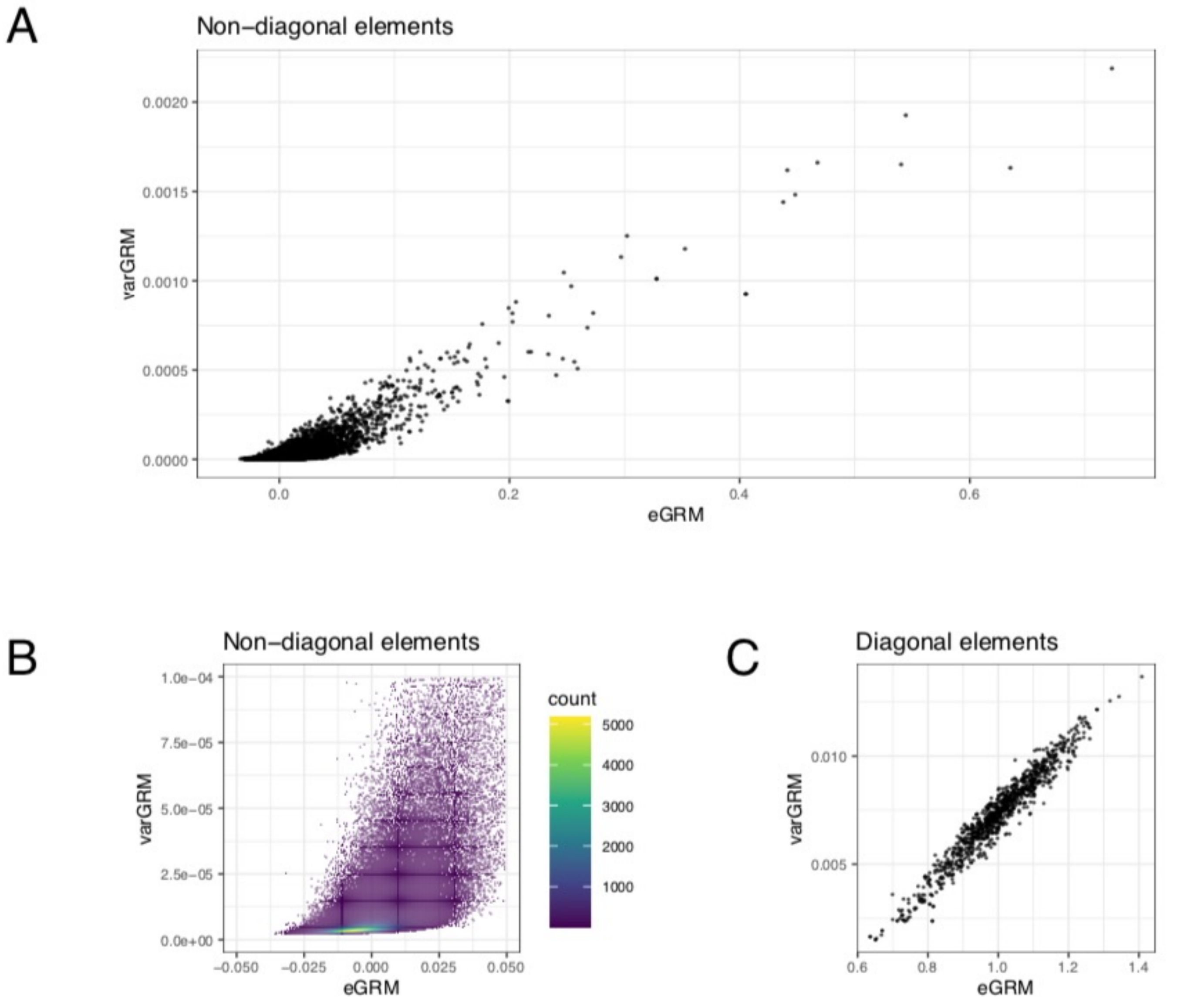


*Figure S8*

**Relationship between eGRM and varGRM values for 1000 samples simulated on a 30Mb chromosome.** **(A)** Scatterplot on non-diagonal elements between eGRM and varGRM. (**B**) a zoom-in of (**A**). (**C**) Scatterplot of the diagonal elements between eGRM and varGRM.
