## Supplemental Methods for "A genealogical estimate of genetic relationships"

**Supplementary Method**

*Diploid GRM and eGRM*

Here, we extend our haploid-based definitions of relatedness to diploid genotypes. We denote the Fraktur letters $\mathfrak{m}$ and $\mathfrak{p}$ as the indexes of the $N/2$ maternal and paternal haplotypes, respectively. In this section we use the notation $X_{\mathfrak{a}}^{\mathfrak{b}}$ to indicate the submatrix of $A$ with rows (i.e., variants) indexed by $\mathfrak{a}$ and columns (i.e., haplotypes) indexed by $\mathfrak{b}$. The diploid genotype matrix is given by $Y = X^{\mathfrak{m}} + X^{\mathfrak{p}}$ and note that,

$$Y_k {Y_k}^T = (X^{\mathfrak{m}} + X^{\mathfrak{p}})_k {(X^{\mathfrak{m}} + X^{\mathfrak{p}})_k}^T = (X^{\mathfrak{m}})_k {(X^{\mathfrak{m}})_k}^T + (X^{\mathfrak{m}})_k {(X^{\mathfrak{p}})_k}^T + (X^{\mathfrak{p}})_k {(X^{\mathfrak{m}})_k}^T + (X^{\mathfrak{p}})_k {(X^{\mathfrak{p}})_k}^T$$

$$= \left(X_k X_k^T\right)_{\mathfrak{m}}^{\mathfrak{m}} + \left(X_k X_k^T\right)_{\mathfrak{m}}^{\mathfrak{p}} + \left(X_k X_k^T\right)_{\mathfrak{p}}^{\mathfrak{m}} + \left(X_k X_k^T\right)_{\mathfrak{p}}^{\mathfrak{p}}$$

And

$$\overline{Y_k} = \overline{(X^{\mathfrak{m}} + X^{\mathfrak{p}})_k} = 2\overline{X_k}$$

The genotype-based GRM is defined as

$$K_{\text{diploid}}(Y) = \frac{1}{M} \sum_{1 \le k \le M} \frac{(Y_k - \overline{Y_k}\mathbf{1})(Y_k - \overline{Y_k}\mathbf{1})^T}{\overline{Y_k}(1 - \overline{Y_k}/2)} = \frac{1}{M} C_{N/2} \left( \sum_{1 \le k \le M} \frac{Y_k {Y_k}^T}{\overline{Y_k}(1 - \overline{Y_k}/2)} \right) C_{N/2})$$

$$= \frac{1}{M} C_{N/2} \left( \sum_{1 \le k \le M} \frac{1}{2\overline{X_k}(1 - \overline{X_k})} \left( \left(X_k X_k^T\right)_{\mathfrak{m}}^{\mathfrak{m}} + \left(X_k X_k^T\right)_{\mathfrak{m}}^{\mathfrak{p}} + \left(X_k X_k^T\right)_{\mathfrak{p}}^{\mathfrak{m}} + \left(X_k X_k^T\right)_{\mathfrak{p}}^{\mathfrak{p}} \right) \right) C_{N/2}$$

$$= \frac{1}{2} \left( \left(K(X)\right)_{\mathfrak{m}}^{\mathfrak{m}} + \left(K(X)\right)_{\mathfrak{m}}^{\mathfrak{p}} + \left(K(X)\right)_{\mathfrak{p}}^{\mathfrak{m}} + \left(K(X)\right)_{\mathfrak{p}}^{\mathfrak{p}} \right).$$

Therefore, the genotype-based eGRM is given by,

$$E\left(K_{\text{diploid}}(Y)\right) = \frac{1}{2} \left( \left(E\left(K(X)\right)\right)_{\mathfrak{m}}^{\mathfrak{m}} + \left(E\left(K(X)\right)\right)_{\mathfrak{m}}^{\mathfrak{p}} + \left(E\left(K(X)\right)\right)_{\mathfrak{p}}^{\mathfrak{m}} + \left(E\left(K(X)\right)\right)_{\mathfrak{p}}^{\mathfrak{p}} \right).$$

This means that starting from a haploid eGRM, if we take the four submatrices indexed by the maternal and paternal chromosomes, half of their summation will be the diploid eGRM.

*Variance of pair-wise relatedness given a genealogy*

Given our probabilistic formulation of relatedness given a genealogy, it is natural to define higher central moments. Here, we define the element-wise $\mathrm{Var}(K(X)|G)$, which captures the expected deviation around the individual entries in the eGRM, which we term varGRM. While the complete variance-covariance matrix of $K(X)$ is $N^2 \times N^2$, our definition reflects solely its $N \times N$ diagonal values, which translates to marginal element-wise variances in the eGRM. By the law of total variance

$$\mathrm{varGRM} \coloneqq \mathrm{Var}(K(X)|M>0) = \mathrm{Var}(E(K(X)|M)|M>0) + E(Var(K(X)|M)|M>0)$$

To simplify this, we first compute the conditional expectation and variance of $K(X)$. Using the fact that whenever $M > 0$, $M(e)|M \sim \mathrm{Binom}\left(M, \frac{\mu(e)}{\mu(G)}\right)$, we have

$$E(K(X)|M) = E\left(\sum_{e \in G} \frac{M(e)}{M} K\big(x(e)\big)\,|M\right)$$

$$= \frac{1}{M} E\left(\sum_{e \in G} M(e) K\big(x(e)\big)\,|M\right) = \frac{1}{M}\sum_{e \in G} M \frac{\mu(e)}{\mu(G)} K\big(x(e)\big) = \sum_{e \in G} \frac{\mu(e)}{\mu(G)} K\big(x(e)\big) = E(K(X)|M>0)$$

Which means the conditional expectation is independent from $M$. Now

$$\mathrm{Var}(K(X)|M) = \mathrm{Var}\left(\sum_{e \in G} \frac{M(e)}{M} K\big(x(e)\big)\,|M\right) = \frac{1}{M^2}\mathrm{Var}\left(\sum_{e \in G} M(e) K\big(x(e)\big)\,|M\right)$$

$$= \frac{1}{M}\mathrm{Var}\left(\sum_{e \in G} M(e) K\big(x(e)\big)\,|M=1\right)$$

The last equation is because a multinomial distribution with $M$ trials is the sum of $M$ i.i.d. multinomial distributions with one trial and same probabilities.

Taken together we have

$$\mathrm{varGRM} \coloneqq \mathrm{Var}(K(X)|M>0) = \mathrm{Var}(E(K(X)|M)|M>0) + E(\mathrm{Var}(K(X)|M)|M>0)$$

$$= \mathrm{Var}(E(K(X)|M>0)|M>0) + E\left(\frac{1}{M}\mathrm{Var}\left(\sum_{e \in G} M(e) K\big(x(e)\big)\,|M=1\right)|M>0\right)$$

$$= 0 + E\left(\frac{1}{M}|M > 0\right) Var\left(\sum_{e \in G} M(e) K\big(x(e)\big)\,|M = 1\right)$$

The expectation $E\left(\frac{1}{M}|M > 0\right)$ is the negative moment of the Poisson distribution and can be approximated numerically. And

$$Var\left(\sum_{e \in G} M(e) K\big(x(e)\big)\,|M = 1\right)$$

$$= E\left(\left(\sum_{e \in G} M(e) K\big(x(e)\big)\right)^2 |M = 1\right) - E^2\left(\sum_{e \in G} M(e) K\big(x(e)\big)\,|M = 1\right)$$

The second term

$$E^2\left(\sum_{e \in G} M(e) K\big(x(e)\big)\,|M = 1\right) = E^2(K(X)|M = 1) = E^2\big(K(X)\big) = \mathrm{eGRM}^2$$

The first term

$$E\left(\left(\sum_{e \in G} M(e) K\big(x(e)\big)\right)^2 |M = 1\right) = E\left(\sum_{e \in G} M(e) K^2\big(x(e)\big)\,|M = 1\right)$$

$$= E\left(\sum_{e \in G} \frac{M(e)}{M} K^2\big(x(e)\big)\right) = \sum_{e \in G} \frac{\mu(e)}{\mu(G)} K^2\big(x(e)\big)$$

The equality in the first line is because when $M = 1$, only one of the branches has $M(e) = 1$, and for any other branch $M(e') = 0$. We now have

$$\mathrm{varGRM} := \mathrm{Var}\big(K(X)\big) = E\left(\frac{1}{M}\right)\left(\sum_{e \in G} \frac{\mu(e)}{\mu(G)} K^2\big(x(e)\big) - \mathrm{eGRM}^2\right)$$

Note that here $K^2\big(x(e)\big)$ is element-wise square, which has to be handled with care

$$\frac{\mu(e)}{\mu(G)} K^2\big(x(e)\big) = \frac{\mu(e)/\mu(G)}{\left[\overline{x(e)}\big(1 - \overline{x(e)}\big)\right]^2}\left(\big(x(e) - \overline{x(e)}\mathbf{1}\big)\big(x - \overline{x(e)}\mathbf{1}\big)^T\right)^2$$

And

$$\left(\big(x(e) - \overline{x(e)}\mathbf{1}\big)\big(x - \overline{x(e)}\mathbf{1}\big)^T\right)^2 = \left(x(e)x(e)^T - \overline{x(e)}\mathbf{1}x(e)^T - \overline{x(e)}x(e)\mathbf{1}^T + \overline{x(e)}^2\mathbf{1}\mathbf{1}^T\right)^2$$

$$= \left(1 - 2\overline{x(e)}\right)^2 x(e)x(e)^T + \left(\overline{x(e)}^2 - 2\overline{x(e)}^3\right)\mathbf{1}x(e)^T + \left(\overline{x(e)}^2 - 2\overline{x(e)}^3\right)x(e)\mathbf{1}^T + \overline{x(e)}^4\mathbf{1}\mathbf{1}^T$$

Computationally, the varGRM is achieved by traversing the ARG in any order, updating a buffer matrix $A$, a buffer vector $\boldsymbol{a}$ and a buffer scalar $a$. So that after traversing we get

$$A = \sum_{e\in G} \frac{\mu(e)/\mu(G)}{\left[\overline{x(e)}\left(1 - \overline{x(e)}\right)\right]^2} \left(1 - 2\overline{x(e)}\right)^2 x(e)x(e)^T$$

$$\boldsymbol{a} = \sum_{e\in G} \frac{\mu(e)/\mu(G)}{\left[\overline{x(e)}\left(1 - \overline{x(e)}\right)\right]^2} \left(\overline{x(e)}^2 - 2\overline{x(e)}^3\right)x(e)$$

$$a = \sum_{e\in G} \frac{\mu(e)/\mu(G)}{\left[\overline{x(e)}\left(1 - \overline{x(e)}\right)\right]^2} \overline{x(e)}^4$$

Finally, we have

$$\mathrm{varGRM} = E\left(\frac{1}{M}\right)\left(A + \mathbf{1}\boldsymbol{a}^{\boldsymbol{T}} + \boldsymbol{a}\mathbf{1}^{\boldsymbol{T}} + a\mathbf{1}\mathbf{1}^{\boldsymbol{T}} - \mathrm{eGRM}^2\right).$$

Even though the practical usage of the concept of varGRM is still to be explored, it provides interesting insights to our understanding of GRM matrices (**Figure S8)**. Note that positive values in the GRM matrix have greater uncertainties than that of negative values, which challenges the possibly common thinking that the sign of GRM is arbitrary. This is because the ancestral allele is represented as 0, while the derived allele is represented as 1 in the genotype matrix, and derived alleles have overall lower frequencies than ancestral alleles. It's also worth noting that the square root of varGRM is at the same order of magnitude as GRM (here the number of variants is ~10,000), which implies that individual elements in the GRM matrix has considerable randomness.

*Average-case runtime complexity*

We denote the expected number of operations to compute the $N \times N$ eGRM given a single genealogical tree as $S(N)$. If the first split from the root divides the tree into two subtrees with samples sizes $x$ and $N - x$, then $S(N)$ can be divided into two parts: 1) the runtime on the two branches from the root $x^2 + (N-x)^2$, because the number of operations to handle each branch is the squared number of its descendants; 2) the time on the two subtrees $S(x) + S(N-x)$. We have

$$S(N) = \sum_{x=1}^{N-1} P(x)\left(x^2 + (N-x)^2 + S(x) + S(N-x)\right)$$

$S(N)$ depends on the distribution of $x$, which is decided by the demography of the population. In a homogeneous population, all allowed topologies of the tree are equally likely. We denote the number of possible topologies of $N$ samples as $T(N)$, then

$$T(N) = \binom{N}{2} T(N-1) = \binom{N}{2}\binom{N-1}{2} T(N-2) = \cdots = \prod_{i=2}^{N} \binom{i}{2} = \frac{N!\,(N-1)!}{2^{N-1}}$$

Note that, there are $\frac{1}{2}\binom{N}{x}$ ways to select $x$ samples to be included in the first subtree ("first" can be defined as, for example, including the first sample). If the topologies within two subtrees are fixed, there are still $\binom{N-2}{x-1}$ possible relative orders of their coalescent events (there are $x-1$ coalescent events in the first subtree and $N-x-1$ in the second, summing up to $N-2$). So we have

$$P(x) = \frac{1}{2}\binom{N}{x}\binom{N-2}{x-1}\frac{T(x)T(N-x)}{T(N)}$$

$$= \frac{N!}{x!\,(N-x)!}\frac{(N-2)!}{(x-1)!\,(N-x-1)!}\frac{x!\,(x-1)!\cdot(N-x)!\,(N-x-1)!}{N!\,(N-1)!} = \frac{1}{N-1}$$

Which means $x$ follows a uniform distribution. So, the average-case time complexity is

$$S(N) = \sum_{x=1}^{N-1} \frac{1}{N-1}\left(x^2 + (N-x)^2 + S(x) + S(N-x)\right) = \frac{2}{N-1}\sum_{i=1}^{N-1} i^2 + \frac{2}{N-1}\sum_{i=1}^{N-1} S(i)$$

Which is equivalent to the recursive equation

$$S(N) = \frac{N}{N-1} S(N-1) + 2(N-1)$$

With boundary condition $S(1) = 1$, The solution is

$$S(N) = 2N^2 + N - 2N\sum_{i=1}^{N}\frac{1}{i} = \Theta(N^2)$$

So, we have proven the average-case time complexity is quadratic to the sample size.
